# History biases reveal novel dissociations between perceptual and metacognitive decision-making

**DOI:** 10.1101/737999

**Authors:** Christopher S.Y. Benwell, Rachael Beyer, Francis Wallington, Robin A. A. Ince

**Affiliations:** Division of Psychology, School of Social Sciences, University of Dundee, Dundee, UK; Institute of Neuroscience and Psychology, University of Glasgow, Glasgow, UK

**Keywords:** Choice history bias, serial dependence, perception, metacognition, confidence

## Abstract

Human decision-making and self-reflection often depend on context and internal biases. For instance, decisions are often influenced by preceding choices, regardless of their relevance. It remains unclear how choice history influences different levels of the decision-making hierarchy. We employed analyses grounded in information and detection theories to estimate the relative strength of perceptual and metacognitive history biases, and to investigate whether they emerge from common/unique mechanisms. Though both perception and metacognition tended to be biased towards previous responses, we observed novel dissociations which challenge normative theories of confidence. Different evidence levels often informed perceptual and metacognitive decisions within observers, and response history distinctly influenced 1^st^ (perceptual) and 2^nd^ (metacognitive) order decision-parameters, with the metacognitive bias likely to be strongest and most prevalent in the general population. We propose that recent choices and subjective confidence represent heuristics which inform 1^st^ and 2^nd^ order decisions in the absence of more relevant evidence.

## Introduction

Human knowledge of the external world and of internal cognitive processes is often biased and incomplete^1–3^. When decisions are made about sensory input (i.e. Is a target present?), we can distinguish between objective accuracy (perceptual sensitivity), and how accurate one is in judging their own performance (metacognitive insight)^4,5^. Metacognitive insight can be quantified by comparing subjective confidence to objective accuracy^4–6^. Though accuracy and confidence usually correlate, metacognitive performance differs widely across individuals^3,7,8^ with important consequences in everyday life. For instance, insight modulates learning, adaptive decision-making, error monitoring and exploration^9–13^. In fact, impaired metacognition is associated with many neuropsychiatric disorders^14^ and sub-clinical symptom dimensions^15^.

Even in healthy individuals, perceptual and metacognitive decisions not only depend on the immediately available evidence, but also on recent experiences and choices. For instance, when similar stimuli are serially presented, perceptual decisions are often biased towards responses and/or stimuli on preceding trials, a phenomenon known as choice history bias^16–21^ or serial dependence^22–27^. Whilst this mechanism may generally be adaptive, because recent experience usually predicts upcoming input, it can also lead to non-veridical decisions^23,28–30^. Interestingly, serial dependence has also been reported for subjective confidence reports^31^, and the level of confidence on the preceding trial has been suggested to modulate perceptual history bias, with repetition more likely when preceding confidence was high^16,17,32^. These reports suggest the existence of an intimate link between perception and metacognition in the formation of history biases. However, the exact nature of this relationship, and the relative strength and source of each bias, remain unclear.

Employing both model-based and non-parametric analyses, we observed history biases in both perceptual responses and ratings of confidence, but we show that the metacognitive history bias is stronger and likely to be most prevalent in the general population. Computational modelling revealed intriguing dissociations between perceptual and metacognitive decision-making parameters. For instance, perceptual choice alternation (disengagement from hysteresis) was associated with increased perceptual sensitivity but reduced metacognitive insight. Overall performance closely matched predictions from recently proposed computational models of decision-making and confidence^33–35^. However, we crucially demonstrate that both perceptual and metacognitive decision criteria are not fixed: they fluctuate from moment-to-moment and are biased by recent choices. Accurate models of subjective confidence must go beyond a normative account to capture sub-optimal metacognitive performance driven by irrelevant factors such as preceding confidence reports.

## Results

### Overall performance exhibited signatures predicted by computational models of decision-making and metacognition

Thirty-seven human observers performed a two-alternative forced choice (2-AFC) orientation discrimination task (Fig. 1A and Methods). Participants judged whether a briefly presented Gabor patch was tilted leftward or rightward of the vertical plane, and reported the level of confidence they felt in their decision (on a scale of 1 – ‘Not confident at all’ to 4 – ‘Highly confident’). The true orientation (and hence task difficulty) was manipulated from trial-to-trial. This design allowed us to test predictions arising from a recently proposed computational model of perceptual decision-making and metacognition based on Bayesian statistical confidence and signal detection theory (SDT), as defined in Figure 1B (and Methods). Briefly, human decisions have been modelled as the comparison of an internal decision variable (DV), representing the evidence in favour of one or other choice in 2-AFC tasks, against a decision criterion (C). Under this model, confidence in the decision is given by the distance of the DV from C^4,5,16,33–35^. When a discrete confidence rating scale is employed, the level of confidence is defined by where the DV falls with respect to the so-called type-2 criteria (*c_1_*, *c_2_*, … *c_N−1)_*), where N indexes the number of possible ratings. A confidence rating of k will follow if the DV falls in the interval (*c_k−1_*, *c_k_*).

**Figure 1:**
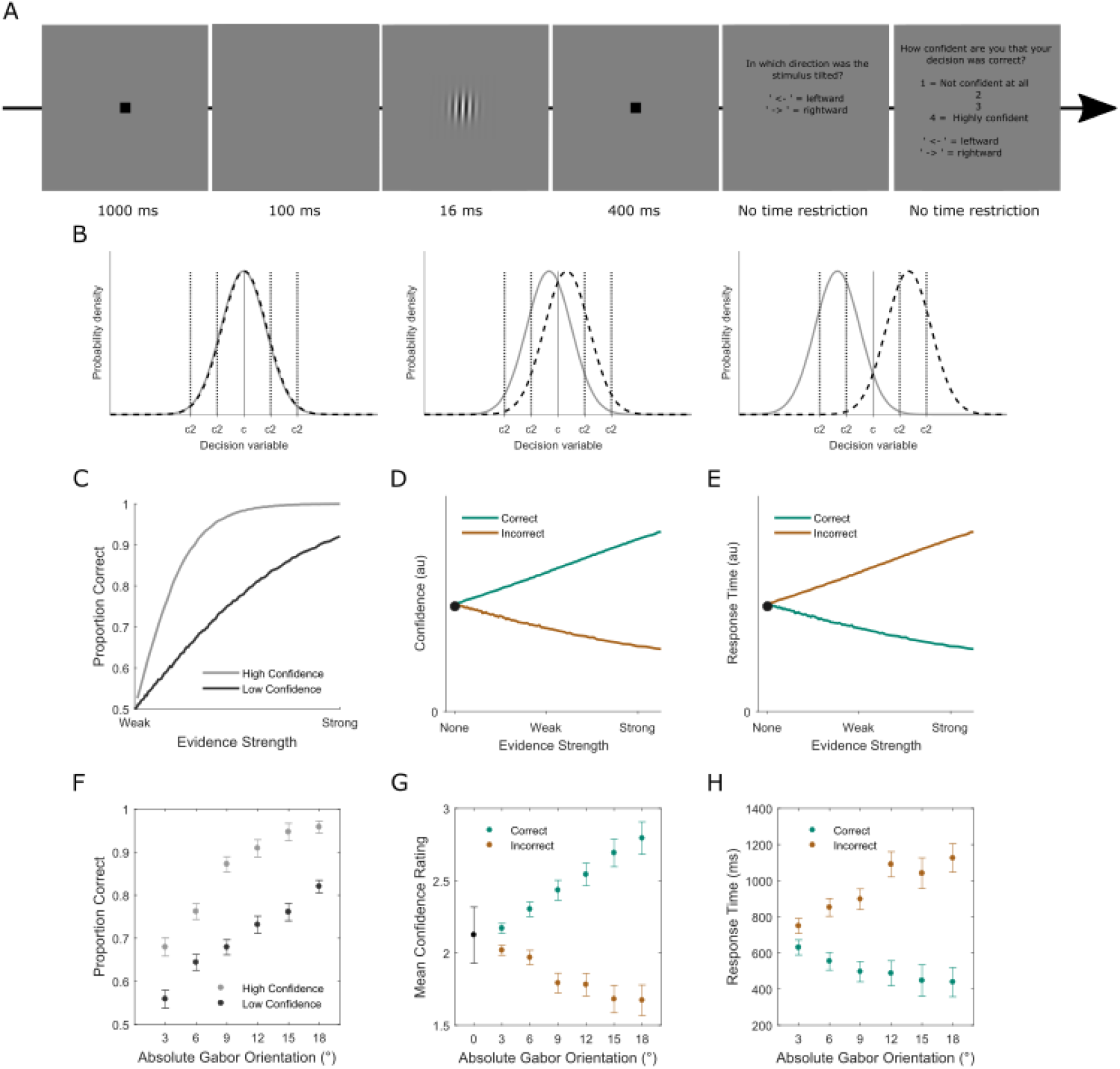
**(A)** Behavioural task. On each trial, a Gabor orientation discrimination judgement was made followed by a confidence report (scale of 1 to 4, where 1 represented “not confident at all” and 4 represented “highly confident”). **(B)** Computational model of decision making and confidence in a 2-AFC task. The probability density functions represent distributions of internal responses (decision variables (DV)) across repeated presentations of the generative stimulus. On each trial, the DV is drawn from one of these distributions and compared with a decision criterion (*c*: solid black vertical line) in order to reach a binary choice. The level of confidence in the choice is then reflected in the absolute distance of the DV from *c*. When a discrete confidence rating scale is employed, the level of reported confidence is defined by where the DV falls with respect to the type-2 criteria (*c_1_*, *c_2_*, … *c_(N−1)_*: dashed vertical black lines), where N indexes the number of possible ratings. A confidence rating of k will be given if the DV falls in the interval (*c_k−1_*, *c_k_*). The relative separation on the x-axis of the two distributions indexes the level of evidence available for the decision. The model is plotted for three levels of overall decision evidence: none (left panel), weak (centre panel) and strong (right panel). **(C)** Model-based prediction of the relationship between decision accuracy and evidence strength as a function of confidence level. **(D)** Predicted relationship between decision confidence and evidence strength as a function of accuracy. **(E)** Predicted relationship between response time and evidence strength as a function of accuracy. These model-based predictions were all confirmed in the data. **(F)** Relationship between decision accuracy and absolute Gabor orientation as a function of confidence level. **(G)** Relationship between decision confidence and absolute Gabor orientation as a function of accuracy. **(H)** Relationship between response time and evidence strength as a function of accuracy.

This model gives rise to several predictions regarding the relationships between stimulus evidence, accuracy and decision confidence^16,34,36–38^: (1) Accuracy should scale with evidence strength (Fig. 1C); (2) Conditioning type-1 performance on high or low confidence ratings should change the slope of the relationship between stimulus evidence and accuracy, with a steeper slope for high relative to low confidence trials (Fig. 1C); (3) Confidence should increase with evidence strength for correct trials, but decrease with evidence strength for incorrect trials (Fig. 1D); (4) Even when there is no veridical evidence in favour of one response or other, confidence should be above 0 (Fig. 1D). These predictions were all confirmed in our data. Accuracy increased as a function of evidence strength but the slope of the stimulus evidence-accuracy relationship was steeper for high relative to low confidence trials (Fig. 1F). Confidence increased with evidence strength for correct trials but decreased with evidence strength for incorrect trials (Fig. 1G). Accordingly, response time decreased as a function of evidence strength for correct trials but increased for incorrect trials (Fig. 1H). Finally, participants reliably reported some level of confidence in decisions even when the Gabor patch was vertically aligned and hence there was no informative evidence (Fig. 1G).

### Choice history bias occurs in both perceptual and metacognitive decisions, but is stronger in metacognition

Next, we investigated the degree to which choice history biases both perceptual and metacognitive responses. Across all trials, no systematic group-level bias in favour of either choice was apparent (t-test of psychometric function (PF) thresholds versus 0°: t(36) = .1497, p = .8818, Bayes Factor (BF_10_) = 0.179) (Fig. 2A). However, group-averaged PFs conditioned on the previous response were shifted towards the previous response (‘left’/’right’ responses were more likely following ‘left’/’right’ responses respectively) despite randomly ordered presentations (Fig. 2A). Post-left PF thresholds were significantly biased away from veridical 0° (t(36) = 3.1295, p = .0035, BF_10_= 10.462), as were post-right PF thresholds, but in the opposite direction (t(36) = −2.5466, p = .0153, BF_10_ = 2.9235). Accordingly, post-left were significantly different to post-right thresholds (t(36) = 4.2498, p < .001, BF_10_ = 177.4). The effect remained significant for trial lags of two (t(36) = 5.9966, p < .001, BF_10_ = 2.3930e+04) and three (t(36) = 5.91, p < .001, BF_10_ = 1.8667e+04) (Fig. 2B). It has recently been suggested that confidence on a given trial modulates the likelihood of the perceptual choice being subsequently repeated^16,32^. However, we did not find any influence of preceding confidence on perceptual history bias, with the bias occurring both when confidence was low versus high on the previous trial (Supplementary Fig. 1).

**Figure 2:**
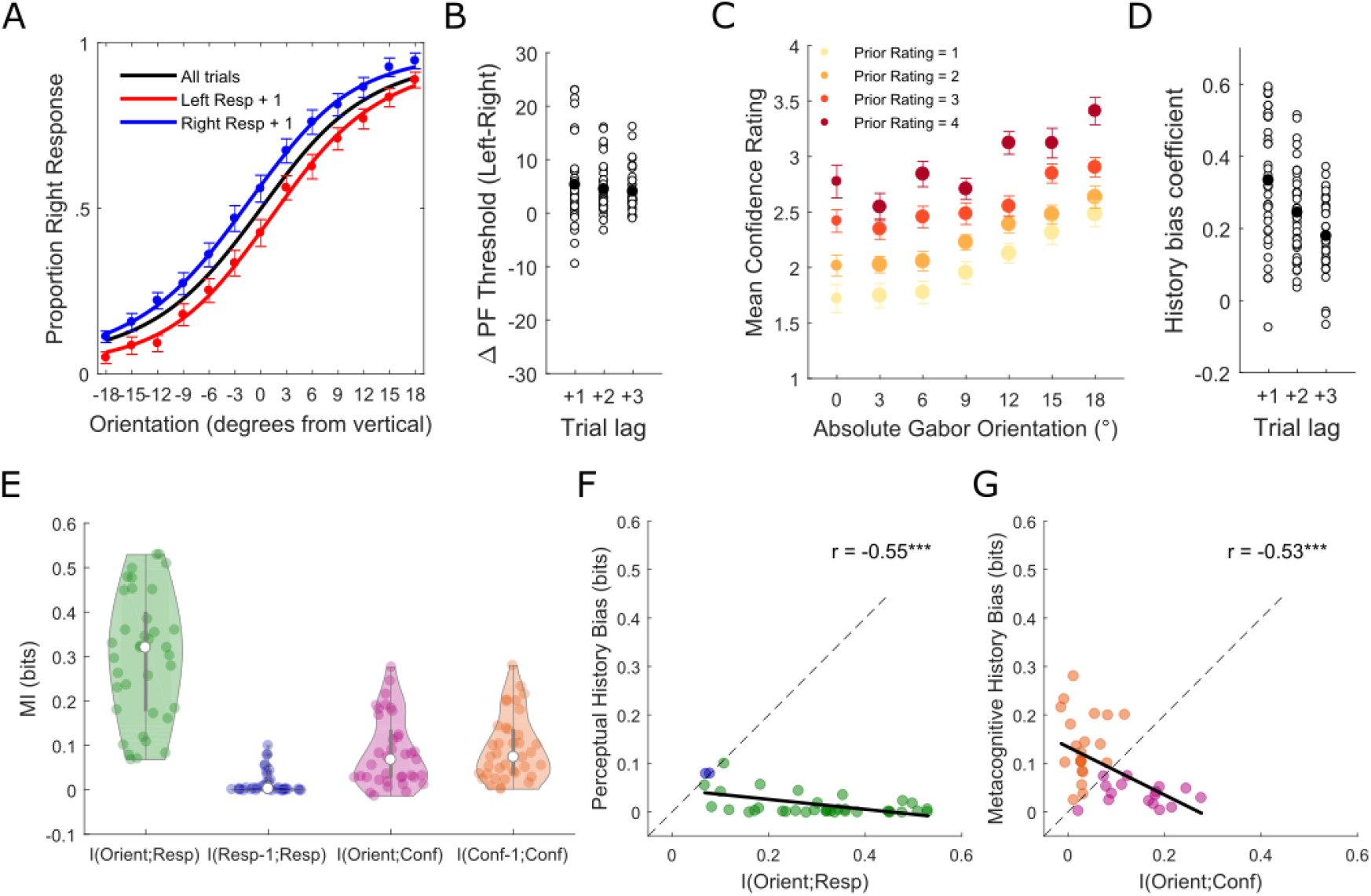
**(a)** Choice history biases perceptual decisions. Group-averaged psychometric functions (PFs) across all trials and conditioned on the previous perceptual choice. **(b)** Scatterplot of single-participant differences in PF threshold between ‘post left choice’ and ‘post right choice’ trials at trial lags of 1, 2 and 3 (black filled dots represents the group means). Positive values index a bias in favour of repeating the previous choice and negative values index a bias in favour of alternating the previous choice. **(c)** Choice history biases confidence ratings. Group-averaged confidence ratings as a function of absolute Gabor orientation on the current trial and rating on the previous trial. The size of the dots indexes the relative number of trials contributing to the group-average as this was not uniform across orientations and previous ratings. **(d)** Scatterplot of single-participant regression coefficients for the linear relationship between confidence on the previous and current trials at lags of 1, 2 and 3. Positive values index that ‘high’/’low’ confidence ratings were more likely following ‘high’/’low’ ratings respectively. All error bars represent within-subject ± standard error (SEM). **(e)** Non-parametric within-participant MI analysis quantified the dependence between evidence presented on each trial (i.e. the orientation of the Gabor) and the perceptual responses/confidence ratings and, on the same effect size scale, the choice history biases in both perceptual responses and confidence ratings. **(f)** The relationship between perceptual choice history bias and the trial-by-trial influence of evidence on the perceptual decision. The influence of evidence was stronger in most participants than the influence of choice history. **(g)** The relationship between metacognitive choice history bias and the trial-by-trial influence of evidence on confidence ratings. There were relatively even sub-groups of participants for whom current evidence dominated confidence judgements vs those for whom choice history dominated confidence judgements. Solid black lines represent least-squares regression slopes. \*\*\**P*<0.001, \*\**P*<0.01, \**P*<0.05, NS *P*>0.05.

Next, we investigated the degree of metacognitive history bias^31^. Confidence increased as a function of absolute orientation (i.e. sensory evidence) but was also shifted towards previous trial ratings (i.e. ‘high’/’low’ were more likely following ‘high’/’low’ ratings respectively) (Fig. 2C). A regression analysis (see Method) confirmed that confidence was positively predicted by ratings on the previous trial across participants (t-test of slopes versus 0: t(36) = 11.7028, p < .001, BF_10_ = 9.3215e+10). The effect remained significant for trial lags of two (t(36) = 11.9084, p < .001, BF_10_ = 1.5082e+11) and three (t(36) = 10.1775, p < .001, BF_10_ = 2.2515e+09) (Fig. 2D).

In order to calculate within-participant significance and estimate population prevalence of the observed biases, we performed additional analyses using mutual information (MI)^39^. MI provides an assumption free measure of dependence with effect sizes on a common meaningful scale (bits) across variables with different characteristics (i.e. different dimensionality and/or number of samples). Hence, we could also quantify and compare how strongly both perceptual and metacognitive responses of each participant were related to the objective evidence at hand versus recent choices. First, we quantified the strength of dependence between stimulus evidence (orientation of the Gabor (Orient)) and both perceptual responses and confidence ratings (Fig. 2E). We then quantified, on the same scale, the choice history biases in both confidence ratings and perceptual responses (see Method for details). Supplementary Figure 2 highlights how the MI measures relate to the model-based bias measures displayed in Fig. 1B and C.

As expected, the highest dependence was found between objective evidence (Orient) and perceptual responses (Resp) (Fig. 2E). Interestingly, this was stronger than the dependence between objective evidence and confidence ratings (t(36) = 11.6448, p < .001, BF_10_ = 8.1307e+10), suggesting sub-optimal metacognitive performance. The confidence history bias was stronger than the perceptual history bias (t(36) = 6.25, p < .001, BF_10_ = 4.9486e+04), and in fact had roughly the same influence on confidence as current trial evidence (t(36) = 0.384, p = .7032, BF_10_ = 0.1894). Statistical inference was performed non-parametrically within individual participants based on 1000 permutations of the data. In our sample, 13/37 participants showed significant perceptual history bias (at p=0.05). Therefore, the population prevalence^40,41^ of perceptual history bias detectable in our experiment is 31.7% [14.6 48.8] (maximum likelihood estimate with 95% bootstrap confidence interval). The majority of those showing significant perceptual history bias tended to repeat their previous responses (N=10), with only 3 tending to alternate^16,19,20^. Across participants perceptual history bias was inversely related to the effect of evidence on perceptual responses within trials (Fig. 2F: Pearson’s r = −0.55, p < .001, BF_10_ = 64.297). However, the influence of evidence was stronger in most participants (MI(Orient;Resp)>MI(Resp-1;Resp), 35/37 participants) than the influence of choice history (MI(Resp-1;Resp)>MI(Orient;Resp), 2/37). 34/37 participants showed significant metacognitive history bias (at p=0.05) which implies a population prevalence of 91.4% [80.1 100]. All participants showing significant metacognitive history bias tended to repeat previous confidence ratings. Across participants metacognitive history bias was inversely related to the effect of evidence on confidence within trials (Fig. 2G: Pearson’s r = −0.53, p < .001, BF_10_ = 29.059), with relatively even sub-groups of participants for whom current evidence dominated confidence judgements (MI(Orient;Conf) > MI(Conf-1;Conf), 16/37), vs those for whom rating history dominated judgements (MI(Conf-1;Conf) > MI(Orient;Conf), 21/37).

### Uncovering the influence of choice history on perceptual and metacognitive decisions with computational behavioural modelling

In order to explore the relationship between perceptual and metacognitive choice history biases, we returned to the decision-making model (defined in Method and Fig. 1B) to formally test which aspects of both perceptual (type-1) and metacognitive (type-2) performance were affected by previous choices. Type-1 performance encompasses traditional measures of perceptual sensitivity (***d’***) and bias (***c***), whereas type-2 performance encompasses measures of metacognitive sensitivity (***meta-d’***) and bias (***meta-c***)^5,42^. ***meta-c*** represents the type-1 ***d’*** value expected to give rise to the observed confidence data under the assumption that the observer has perfect metacognitive insight (i.e. ***d’*** = ***meta-c*** when confidence is always high when correct and low when incorrect). To quantify metacognitive insight/efficiency (or in other words how much of the information present in the type-1 performance participants make use of in their type-2 decisions), we can subtract ***d’*** from ***meta-d’***. If ***meta-d’ -d’*** ≠ 0, then confidence ratings are either more (positive) or less (negative) sensitive to the task-related evidence than the perceptual responses. The metacognitive criteria (***meta-c***) index the tendency to give high/low confidence ratings regardless of evidence (metacognitive response bias). Their absolute distance from type-1 ***c*** (**|*meta-c - c*|**) represents the level of evidence needed to increase confidence ratings from low to high^43^. Unlike type-1 ***c***, ***meta-c*** values are calculated separately for each possible perceptual response (‘left’/‘right’ orientation judgements). Additionally, there are N-1 ***meta-c*** for each response, where N indexes the number of possible ratings (4 in the current experiment). In order to simplify the analysis, we averaged over the 3 **|*meta-c - c*|** values for each response (‘left’/‘right’) separately (see Methods).

First, we assessed whether overall metacognitive sensitivity (***meta-d’***) systematically deviated from perceptual sensitivity (***d’***). Across orientations, confidence judgements were less reflective of the evidence than perceptual judgements, with mean ***meta-d’*** being lower than mean ***d’*** (compare Fig. 3A and 3B). A repeated-measures analysis of variance (ANOVA: 2 (sensitivity measure: ***d’***, ***meta-d’***) × 6 (absolute orientation (evidence): 3°,6°,9°,12°,15°,18°)) revealed that ***meta-d***’ was significantly lower than ***d’*** (main effect: F(1,36) = 58.818, p < .001) and the difference increased as a function of orientation (Fig. 3C) (interaction: F(5,180) = 13.614, p < .001). Hence, participants were generally unable to make use of all information available for perceptual judgements when estimating their confidence (suboptimal metacognition), in line with the MI results. To investigate the influence of previous perceptual choices, we calculated the type-1 and type-2 model parameters^42^ separately for ‘post-left’ and ‘post-right’ decision trials across each level of evidence strength. We then performed repeated-measures ANOVAs (2 (previous choice: ‘left’/‘right’) × 6 (absolute orientation: 3°,6°,9°,12°,15°,18°)) for each parameter.

**Figure 3:**
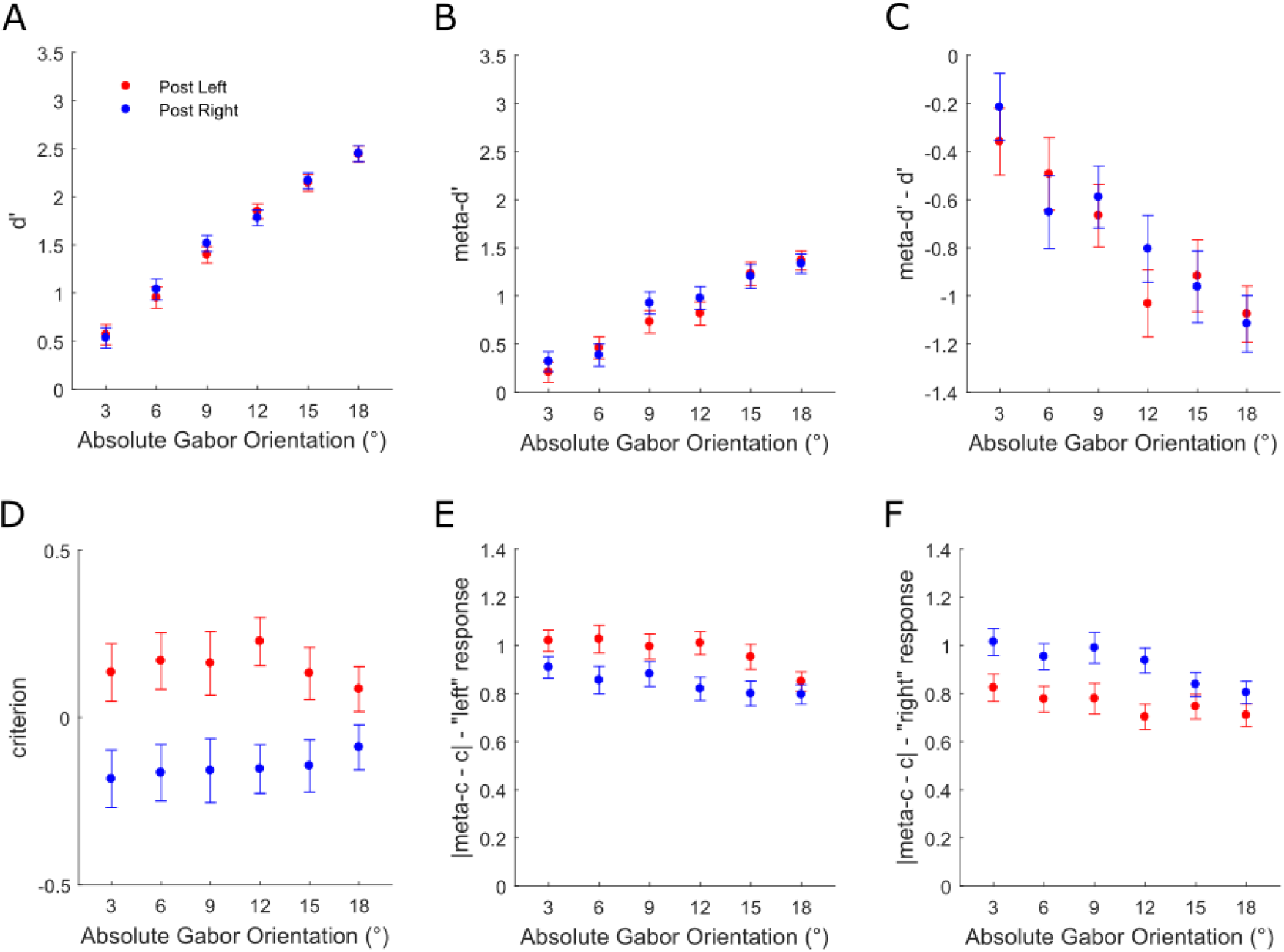
Modelling the influence of perceptual decisions on subsequent perceptual and metacognitive performance (see Method and refs 5,34 for details). **(a)** Group-averaged ***d’*** as a function of absolute Gabor orientation and perceptual choice on the previous trial. **(b)** Group-averaged ***meta-d’*** as a function of absolute Gabor orientation and perceptual choice on the previous trial. **(c)** Group-averaged ***meta-d’ - d’*** as a function of absolute Gabor orientation and perceptual choice on the previous trial. **(d)** Group-averaged ***c*** as a function of absolute Gabor orientation and perceptual choice on the previous trial. **(e)** Group-averaged **|*meta-c - c*|** for ‘left’ responses as a function of absolute Gabor orientation and perceptual choice on the previous trial. **(f)** Group-averaged **|*meta-c - c*|** for ‘right’ responses as a function of absolute Gabor orientation and perceptual choice on the previous trial. All error bars represent within-subject ± standard error (SEM).

Previous perceptual choice did not influence either perceptual or metacognitive sensitivity (Fig. 3A-C), neither ***d’***, ***meta-d’*** nor ***meta-d’–d’*** (F-values ≤ 1.086, p-values ≥ .37). However, type-1 ***c*** was biased towards the previous perceptual choice across all orientations (Fig. 3D: main effect: F(1,36) = 20.344, p < .001; interaction: F(5,180) = 1.619, p = .157), in line with the PF analysis. Metacognitive criteria (**|*meta-c - c*|**) were biased in a response dependent manner (Fig. 3E-F). When participants responded ‘left’, they displayed higher meta-criteria when they had also responded ‘left’ on the previous trial (repetition) compared to when they had responded ‘right’ (alternation) (main effect: F(1,36) = 12.983, p < .001; interaction: F(5,180) = 1.603, p = .162). Accordingly, when participants responded ‘right’, they displayed higher meta-criteria when they had responded ‘right’ on the previous trial (repetition) compared to when they had responded ‘left’ (alternation) (main effect: F(1,36) = 14.52, p < .001; interaction: F(5,180) = 2.427, p = .037). The interaction term in the ‘right’ response analysis was driven by the effect not being significant for the two largest orientations. The effect was significant for 3°, 6°, 9°, 12° (t-values ≥3.164, p-values ≤.003, BF_10_ ≥ 11.352) but not 15° (t(36) = 1.775, p = .084, BF_10_ = 0.731) nor 6° (t(36) = 1.972, p = .056, BF_10_ = 1). In sum, perceptual choices influenced decision criteria for both perceptual and metacognitive subsequent choices.

To investigate the influence of the previous metacognitive choice, we performed the same analysis but this time for ‘post-high’ and ‘post-low’ confidence trials (two bins split as evenly as possible within each participant: see Methods). Previous confidence did not influence perceptual sensitivity (Fig. 4A: ***d’*** main effect: F(1,36) = .076, p = .784; interaction: F(5,180) = .162, p = .976), but it did influence subsequent metacognitive sensitivity (Fig. 4B: ***meta-d’*** (main effect: F(1,36) = 48.972, p < .001; interaction: F(5,180) = 4.617, p = .001)) and metacognitive efficiency (Fig. 4C: ***meta-d’–d’*** main effect: F(1,36) = 33.194, p < .001; interaction: F(5,180) = 2.375, p = .041). The interaction terms in both the metacognitive sensitivity (***meta-d’***) and efficiency (***meta-d’–d’***) analyses were driven by the effect increasing as a function of orientation (Fig. 4B-C). For metacognitive sensitivity, follow-up t-tests showed that the effect was significant for orientations of 3°, 9°, 12°, 15°, 18° (t-values ≥2.413, p-values ≤.021, BF_10_ ≥ 2.239) but not for 6° (t(36) = .457, p = .65, BF_10_ = 0.195). For metacognitive efficiency, the effect was significant for 9°, 12°, 15°, 18° (t-values ≥3.368, p-values ≤.002, BF_10_ ≥ 18.449) but not for 3° (t(36) = 1.737, p = .091, BF_10_ = 0.689) nor 6° (t(36) = .18, p = .858, BF_10_ = 0.179).

**Figure 4:**
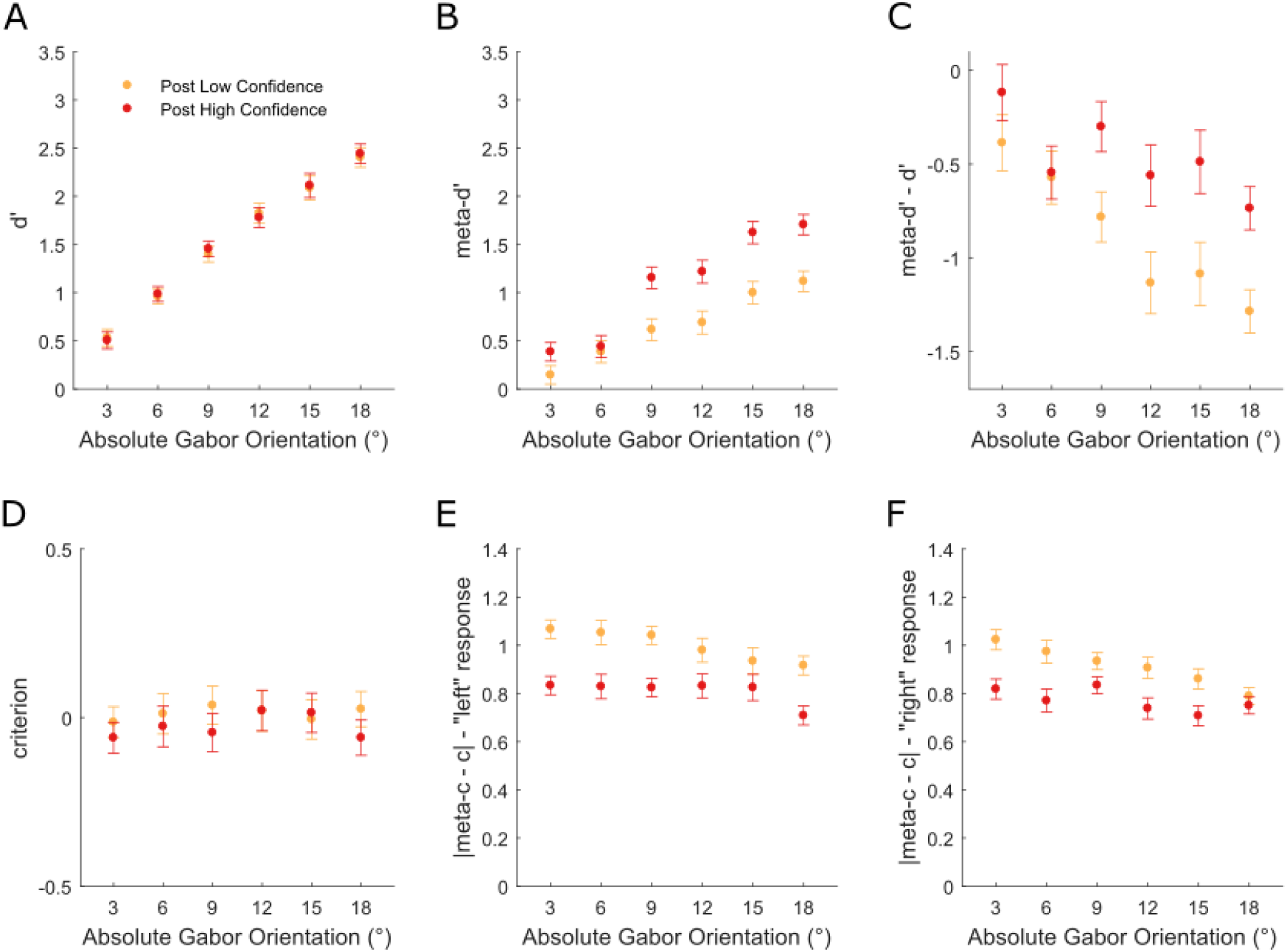
Modelling the influence of metacognitive decisions (confidence ratings) on subsequent perceptual and metacognitive performance. **(a)** Group-averaged ***d’*** as a function of absolute Gabor orientation and confidence on the previous trial. **(b)** Group-averaged ***meta-d’*** as a function of absolute Gabor orientation and confidence on the previous trial. **(c)** Group-averaged ***meta-d’ - d’*** as a function of absolute Gabor orientation and confidence on the previous trial. **(d)** Group-averaged ***c*** as a function of absolute Gabor orientation and confidence on the previous trial. **(e)** Group-averaged **|*meta-c - c*|** for ‘left’ responses as a function of absolute Gabor orientation and confidence on the previous trial. **(f)** Group-averaged **|*meta-c - c*|** for ‘right’ responses as a function of absolute Gabor orientation and confidence on the previous trial. All error bars represent within-subject ± standard error (SEM).

In contrast to the perceptual history bias, type-1 ***c*** was not influenced by confidence on the previous trial (Fig. 4D: main effect: F(1,36) = 1.419, p = .241; interaction: F(5,180) = .645, p = .666). However, **|*meta-c - c*|** were significantly reduced following ‘high’ relative to ‘low’ confidence responses, both for ‘left’ (Fig. 4E) (main effect: F(1,36) = 43.086, p < .001; interaction: F(5,180) = 1.481, p = .198) and ‘right’ responses (Fig. 4F) (main effect: F(1,36) = 31.366, p < .001; interaction: F(5,180) = 3.025, p = .012), indicating that ‘high’/’low’ confidence ratings were more likely following ‘high’/’low’ ratings respectively. The interaction term in the ‘right’ response analysis was driven by the previous rating effect decreasing as a function of orientation (Fig. 4F: linear contrast F(1,36) = 6.771, p = .013). Follow-up t-tests showed that the effect was significant for orientations of 3°, 6°, 9°, 12°, 15° (t-values ≥ 2.739, p-values ≤ .01, BF_10_ ≥ 4.37) but not for 18° (t(36) = 1.066, p = .203, BF_10_ = 0.299). Hence, metacognitive choice history influenced all aspects of metacognitive performance (sensitivity, efficiency and bias), but did not influence perceptual sensitivity nor bias.

### Choice alternation is associated with increased perceptual sensitivity but reduced metacognitive insight

Next, we investigated directly whether repeating (versus alternating) the previous choice was associated with changes in either perceptual and/or metacognitive performance. Figure 5 plots metacognitive history bias effects separately for ‘repetition’ (Fig. 5A) and ‘alternation’ trials (Fig. 5B). For both, confidence increased as a function of orientation but also tended to be shifted towards previous ratings. Confidence was positively predicted by previous ratings for both repetition (t(36) = 11.88, p < .001, BF_10_ = 1.4145e+11) and alternation trials (t(36) = 6.6953, p < .001, BF_10_ = 1.7697e+05). However, the effect was significantly stronger for repetition trials (t(36) = 5.0343, p < .001, BF_10_ = 1.5439e+03). Intriguingly, computational modelling revealed a novel dissociation of perceptual and metacognitive sensitivity induced by disengagement from choice hysteresis. When participants alternated from their previous choice, they were more likely to be correct than when they repeated (Fig. 5C: ***d’*** main effect: F(1,36) = 68.841, p < .001; interaction: F(5,180) = 2.763, p = .02). The effect was significant at all orientations (t-values ≥ 2.709, p-values ≤ .01, BF_10_ ≥ 4.1) but increased as a function of orientation (linear contrast: F(1,36) = 13.284, p = .001). However, this improvement in perceptual sensitivity for alternation trials was not reflected in metacognitive sensitivity (Fig. 5D: ***meta-d’*** (main effect: F(1,36) = 1.311, p = .26; interaction: F(5,180) = .932, p = .462)). Hence, objective accuracy increased for alternation relative to repetition trials whereas metacognitive efficiency decreased (Fig. 5E: ***meta-d’–d’*** main effect: F(1,36) = 27.262, p < .001; interaction: F(5,180) = 2.321, p = .045). The ***meta-d’–d’*** effect was significant at orientations of 6°, 9°, 12°, 15°, 18° (t-values ≤ −2.2, p-values ≤ .033, BF_10_ ≥ 1.552), but not 3° (t(36) = −1.303, p = .201, BF_10_ = 0.385), and increased as a function of orientation (F(1,36) = 9.899, p = .003).

**Figure 5:**
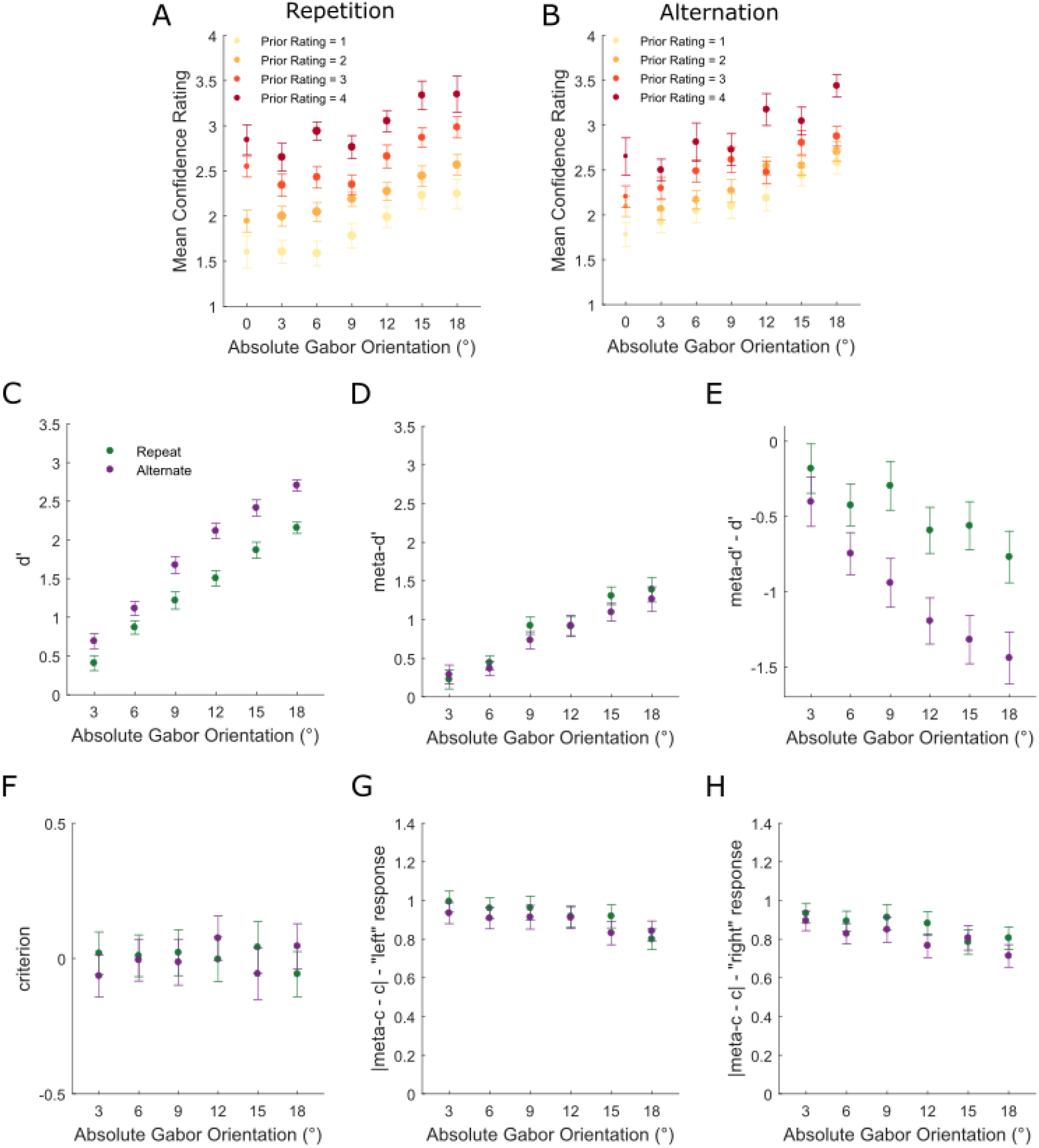
Choice history bias in confidence ratings as a function of perceptual choice hysteresis. **(a)** Group-averaged confidence ratings as a function of absolute Gabor orientation and rating on the previous trial for perceptual choice repetition trials. The size of the dots indexes the relative number of trials contributing to the group-average as this was not uniform across orientations and previous ratings. **(b)** Group-averaged confidence ratings as a function of absolute Gabor orientation and rating on the previous trial for perceptual choice alternation trials. **(c)** Group-averaged ***d’*** as a function of absolute Gabor orientation and perceptual choice relative to previous choice. **(d)** Group-averaged ***meta-d’*** as a function of absolute Gabor orientation and perceptual choice relative to previous choice. **(e)** Group-averaged ***meta-d’ - d’*** as a function of absolute Gabor orientation and perceptual choice relative to previous choice. **(f)** Group-averaged ***c*** as a function of absolute Gabor orientation and perceptual choice relative to previous choice. **(g)** Group-averaged **|*meta-c - c*|** for ‘left’ responses as a function of absolute Gabor orientation perceptual choice relative to previous choice. **(h)** Group-averaged **|*meta-c - c*|** for ‘right’ responses as a function of absolute Gabor orientation and perceptual choice relative to previous choice. All error bars represent within-subject ± standard error (SEM).

Choice hysteresis did not influence either perceptual or metacognitive decision criteria (Fig. 5F-H: F-values ≤ 1.685, p-values ≥ .14). Overall, participants lacked full metacognitive insight into the increased likelihood of being correct when they alternated from their previous perceptual response.

### Choice history biases are associated with reduced perceptual and metacognitive sensitivity, but not reduced metacognitive insight, across participants

Finally, we investigated the correlation between perceptual and metacognitive history biases (Fig. 6A), and whether they contribute to suboptimal perceptual and metacognitive sensitivity, across participants. The strength of perceptual bias did not predict the strength of metacognitive bias (Pearson’s r = 0.1072, p = .5278, BF_10_ = 0.156). Metacognitive history bias was stronger in most participants (MI(Conf-1;Conf)>MI(Resp-1;Resp), 36/37 participants) than perceptual history bias (MI(Resp-1;Resp)>MI(Conf-1;Conf), 1/37 participants).

**Figure 6:**
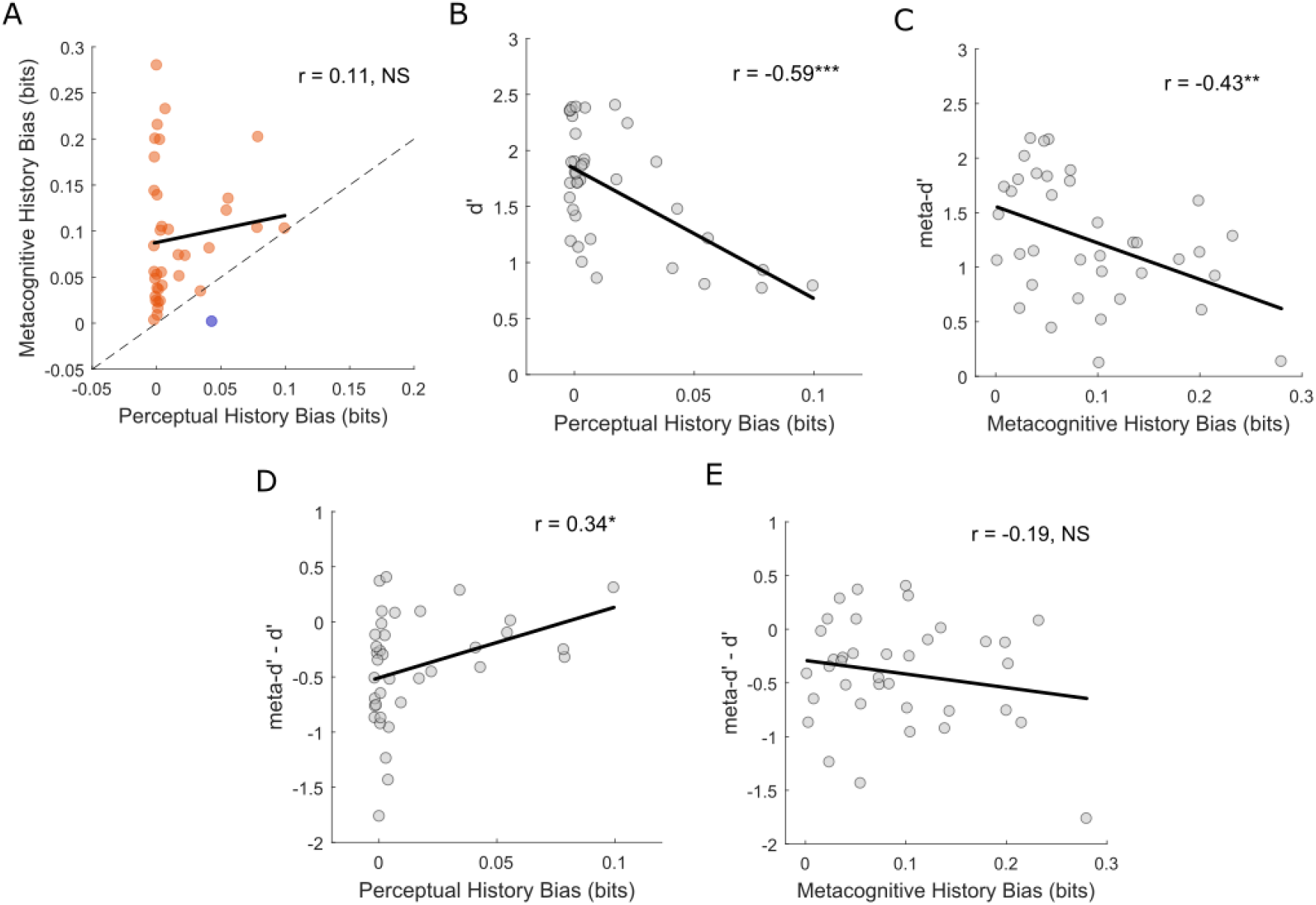
Between-subject Pearson correlations. **(a)** Relationship between perceptual and metacognitive choice history biases. Metacognitive history bias was stronger in most participants (MI(Conf-1;Conf)>MI(Resp-1;Resp), 36/37 participants) than perceptual history bias (MI(Resp-1;Resp)>MI(Conf-1;Conf), 1/37 participants). **(b)** Relationship between perceptual choice history bias and perceptual sensitivity (***d’***). **(c)** Relationship between metacognitive choice history bias and metacognitive sensitivity (***meta-d’***). **(d)** Relationship between perceptual choice history bias and metacognitive efficiency (***meta-d’ - d’***). **(e)** Relationship between metacognitive choice history bias and metacognitive efficiency (***meta-d’ - d’***). Solid black lines represent least-squares regression slopes. \*\*\**P*<0.001, \*\**P*<0.01, \**P*<0.05, NS *P*>0.05.

History biases have previously been linked to reduced perceptual^19^ and metacognitive sensitivity^31^, and we replicated these findings here. The perceptual bias was inversely related to perceptual sensitivity (***d’***) (Fig. 6B: r = −0.5877, p < .001, BF_10_ = 179.836) and the metacognitive bias was inversely related to metacognitive sensitivity (***meta-d’***) (Fig. 6C: r = −0.4315, p = .0077, BF_10_ = 4.353). Perceptual history bias was not significantly associated with metacognitive sensitivity (***meta-d’***) and metacognitive history bias was not significantly associated with perceptual sensitivity (***d’***) (Supplementary Fig. 3).

Previously, a negative correlation was found between metacognitive history bias and metacognitive sensitivity (as quantified by the area under the Type-2 receiver operating characteristic (ROC) curve (Type-2 AUC))^31^. However, neither type-2 AUC nor ***meta-d’*** account for type-1 performance, and hence do not represent pure measures of metacognitive insight/efficiency^5,42^. Therefore, to establish the relationships between perceptual and metacognitive history biases and metacognitive insight, we correlated both with metacognitive efficiency (***meta-d’–d’***). A weak positive correlation was found between perceptual history bias and metacognitive efficiency (Fig. 6D: r = 0.3437, p = .0373, BF_10_ = 1.101). The BF_10_ did not provide strong evidence for the alternative hypothesis therefore we do not interpret this further. However, a one-tailed analysis to test for a negative relationship revealed strong evidence for the null hypothesis (BF_10_ = 0.07). A non-significant negative relationship was observed between metacognitive history bias and metacognitive efficiency (Fig. 6E: r = −0.1852, p = .2726, BF_10_ = 0.233). Hence, when the contribution of type-1 performance to absolute metacognitive sensitivity was factored out, history biases were not significantly associated with reduced metacognitive insight across participants. Note that similar results were found using a ratio measure of metacognitive efficiency/insight (Supplementary Fig. 4).

Using MI to quantify history biases eliminates information about the bias direction (i.e. ‘Repeater’ versus ‘Alternator’). For the sake of completeness, the same correlation analyses using metrics which retain the bias direction are reported in Supplementary Figure 5.

## Discussion

Human decisions are often influenced by sources other than the relevant information^1,2,44,45^. Understanding sub-optimal decision-making represents a fundamental enterprise in modern psychology and neuroscience^46^. In line with previous studies, we show that choice history represents a source of task-irrelevant choice variability, both for perceptual decisions^16–21,27^ and confidence reports^31^. Most participants displayed positive history biases: they were more likely to repeat perceptual decisions and confidence ratings even though stimuli were presented in a random order and hence previous choices were of no relevance. Crucially, we quantified both perceptual and metacognitive history biases on a common effect size scale (using MI) and estimated single-subject significance and population prevalence of the respective effects. Additionally, by employing computational modelling of perceptual decisions and confidence ratings, we were able to uncover latent parameters which are influenced by choice history at different levels of the decision-making hierarchy. Across participants, perceptual and metacognitive history biases did not correlate with each other, but were independently associated with reduced perceptual and metacognitive sensitivity, whereas neither bias predicted metacognitive insight. We show for the first time that the perceptual and metacognitive biases influence distinct type-1 (perceptual) and type-2 (metacognitive) aspects of decision-making, and the metacognitive bias is stronger and likely to be more prevalent in the general population. These observations are of fundamental relevance for contemporary models of decision-making and confidence, suggesting that recent confidence represents a mental shortcut (heuristic) which informs self-reflection when more relevant information is unavailable.

A normative model posits that confidence computations reflect the probability of being correct in a statistically optimal manner^33,34,36,47^, in line with suggestions that the computation of confidence arises from the same neural processes as the decision itself^48–51^. The normative model relates confidence to the available evidence through a conditional Bayesian posterior probability^17,36,44^ and several of the model predictions were met in the current dataset using explicit confidence ratings of visual discrimination performance (see Fig 1B-H). This suggests that subjective confidence is to some extent consistent with normative statistical principles, though it should be noted that a first-order normative model is not the only model which gives rise to such predictions^38^. However, the influence of choice history on confidence ratings (see Fig 2C-G) shows that the normative model alone cannot fully account for subjective confidence. Rather, the normative computation may be one of several determinants of confidence^34^, and differential weighting of these determinants may explain individual differences in the degree of metacognitive history bias and overall metacognitive sensitivity. Other factors which have been suggested to influence confidence judgements include context^52^, social pressure^12,53^, attention^54,55^ and fatigue^56^. Our approach allowed us to quantify and compare the degree to which confidence judgements were driven by objective evidence versus preceding confidence ratings. Surprisingly, we found a relatively even split of participants for whom the objective evidence most strongly influenced confidence versus participants for whom previous ratings were a stronger influence (Fig. 2G). In contrast, all but one participant showed a stronger influence of objective evidence on perceptual choices than the influence of previous choices (Fig. 2F). Metacognitive judgements are thus more susceptible to bias from extraneous factors than perceptual decisions, an observation which may be of practical relevance in terms of learning, error monitoring and psychological well-being^9–13^. Further research may investigate whether primarily ‘history’ versus ‘evidence’ based metacognitive styles meaningfully predict differences in influential traits such as cognitive flexibility, personality and/or psychiatric symptomology.

The current results align with models positing that confidence judgements arise from processes which are to some degree dissociable from the decision process itself^5,38,57^, with distinct neural implementations and independent influences. Evidence supporting such a dissociation has come from neuroimaging^7,8,35,58–64^, psychophysics^32,65,66^, brain stimulation^67,68^ and clinical^14,69,70^ studies. Several aspects of our findings accord with a ‘second-order’ computation of confidence. First, participants were generally unable to make use of all of the information available for their perceptual decisions when rating confidence, which indicates ‘noise’ in the metacognitive system and sub-optimal insight^71,72^. Second, perceptual and metacognitive history biases were uncorrelated across participants (Fig. 6A) and impacted on distinct latent decision-making parameters (Figs. 3+4). For instance, type-2 (metacognitive) decision criteria were modulated as a function of prior confidence ratings independently of the type-1 (perceptual) criteria (Fig. 4D-F), and alternating from choice hysteresis was associated with increased perceptual sensitivity but reduced metacognitive insight (Fig 5A-C). This dissociation when disengaging from choice hysteresis, reported here for the first time, adds to previous reports suggesting that accuracy and confidence can be un-coupled even in healthy participants^55,68,71^. Thus, confidence computations must operate, at least partly, on an axis which is dissociable from type-1 decisions^43^. We did find evidence for some level of interaction between perceptual and metacognitive history biases. The metacognitive bias was strongest for trials in which the perceptual choice was repeated, though it remained significant also for alternation trials (Fig. 5A-B). This suggests that some level of ‘shared’ hysteresis occurs across both systems. However, in contrast to previous findings^16,17,32^, preceding confidence had no influence on the likelihood of the perceptual choice being repeated. Subtle but important differences in experimental designs may explain this discrepancy (see Supplementary Figure 1).

Why might perceptual and metacognitive decision processes be dissociable? One possibility is that the nature of everyday decision-making renders the utilisation of all type-1 information for metacognitive reflection either impossible or unnecessary^71^. As is known for decision-making, metacognitive judgements might rely partly on heuristics and simplifications which result in systematic biases under particular conditions, including lab-based tasks with high levels of uncertainty^8,71,73–75^. In natural settings, it may generally be advantageous to assume statistical regularity of environmental stimuli and to default to this model/heuristic under conditions of high uncertainty^27,28^. If the metacognitive system has less access to objective evidence than the perceptual system, then stronger history biases of confidence ratings are likely to occur. Indeed, here confidence ratings were less sensitive to the objective evidence than perceptual choices and were also more strongly biased by previous ratings.

The mechanisms underlying history biases remain unclear, though neural signatures encoding previous perceptual choices have been identified across various sensory, associative and motor brain regions^26,76–78^. Recent studies have investigated perceptual history bias within the context of computational models of decision-making. The drift-diffusion model (DDM)^79^ represents an extension of classic SDT incorporating single-trial dynamics of evidence accumulation. Under this model, biasing of the type-1 criterion by previous choices (Fig 3D) could occur due to asymmetry in either the starting point or drift rate of the evidence accumulation process. Urai et al.,^20^ show compelling evidence across six tasks that drift rate bias provides the best account, in line with persistence of decisional weights over time/trials^18,27^, an interpretation which is fully in line with our results. An intriguing avenue for further research would be to model the temporal dynamics of both type-1 and type-2 decisions^57^ in order to ascertain the mechanism(s) of history induced type-2 criterion shifts (Fig 4E-F). Additionally, by combining such an approach with functional neuroimaging^35,49^, neural correlates of model parameters may reveal the neural implementations underlying both perceptual and metacognitive choices themselves, along with history biases.

To our knowledge, this study is the first to report estimates of the population prevalence of both perceptual and metacognitive choice history biases. We employed information theoretic statistics to quantify aspects of decision-making within individual participants on a common effect size scale. These measures also enable computationally efficient non-parametric within-participant inference. This novel approach could be widely applied to different questions in studies of decision-making. We found that metacognitive history bias was significant in almost all of our sample (34/37), allowing us to infer an estimate of the population prevalence of 91.5% [80.1 100] (maximum likelihood with 95% bootstrap confidence interval). That is, we can expect that at least 80% of the general population would have an effect detectable with our experimental design (i.e. statistically significant at p=0.05 from 416 trials). The perceptual history bias was significant at the group-level, but was only significant in 13/37 of our sample, yielding a population prevalence estimate of 31.7% [14.6 48.8]. Statistical inference in psychology traditionally focusses on population mean effects, but we argue that it is crucial to determine the degree to which the effects can be reliably observed within individuals, and the prevalence of these effects in the population.

The extent to which these biases negatively influence everyday decisions remains unclear, though repeating previous choices in situations of uncertainty may serve to preserve neural resources associated with choice alternation and to maintain self-consistency^8^. Indeed, activation of a specific cortical network involving inferior frontal cortex and the subthalamic nucleus during the decision process is associated with disengagement from choice hysteresis^80^. This suggests that switching choices under conditions of uncertainty comes at a computational cost. It is interesting to speculate that engagement of this network might improve performance but not subjective confidence in the choice, thereby explaining the lack of metacognitive insight our participants displayed, despite increased perceptual sensitivity, when alternating from their previous choice (Fig. 5A-C). Furthermore, the drive for hysteresis/self-consistency may induce uncertainty when choices are switched and hence distort metacognitive judgements.

It is possible that such biases could have negative implications in circumstances where significant decisions must be made under conditions of high uncertainty (i.e. security scanning, medical imaging^81^). Furthermore, miscalibrated metacognitive judgement (systematic under- or over-confidence) is likely to impact on learning, adaptive decision-making and mental health^9–15^, and may be compounded by history and confirmation biases. The development of behavioural and/or pharmacological techniques to reduce such biases can help to optimise accurate decision-making and self-reflection.

## Method

### Participants

Forty-three healthy human observers participated in the study. All reported normal or corrected-to-normal vision. Due to poor psychophysical performance (explained in the *Data Exclusion* section), 6 participants were excluded from the analysis, leaving a total number of 37 participants (26 female/11 male aged from 18 to 38 years (*M* = 25.23, *SD* = 3.95)). The study was approved by the Ethics Committee of the College of Science and Engineering at the University of Glasgow and all participants gave their informed consent. No monetary reward was given to participants for taking part, though undergraduate students could receive course credits for their participation.

### Stimuli and task

The stimuli were Gabor patches (windowed sine wave gratings: 96 × 96 pixels (2.54 × 2.45 cm)) presented at the centre of the screen. The Gabor patches had a peak contrast of 100% Michelson, a spatial frequency of 3.7 cycles per degree and a 0.3° standard deviation Gaussian contrast envelope. At a viewing distance of 57 cm (fixed using a chinrest), Gabor patches subtended 2.55° of visual angle. On each trial, the stimulus would appear at a random angle that ranged from −18° to +18° relative to vertical at intervals of 3° (including 0°). The monitor used to present the stimuli had a display refresh rate of 60Hz and screen resolution of 1920 × 1080 pixels. The software used to implement the task was E-prime 2.0 and participants made responses using a QWERTY keyboard. Each trial began with a fixation point displayed at the centre of the screen for 1000ms (see Fig. 1A). Following this, a Gabor patch appeared at a random orientation in the centre of the screen for a duration of 16ms. After the stimulus disappeared, the participant viewed the fixation point for 400ms, before being instructed to indicate whether they perceived the top of the Gabor patch to be tilted in a “leftward” or “rightward” direction relative to vertical (2-alternative forced choice), by responding with the left and right arrow keys respectively. Participants were not informed as to the accuracy of their choice and no time limit was enforced. Immediately after responding, participants were presented with a second decision regarding their confidence about the perceptual choice they had just made. Participants were asked to rate their confidence on a scale of 1 to 4, where 1 represented “not confident at all” and 4 represented “highly confident”, using the corresponding digit keys on the keyboard. Immediately after making this response the central fixation point reappeared indicating the beginning of the next trial. A short practice block (12 trials), including only the most extreme angles (−18°, +18°) and with accuracy feedback on each trial, was performed in order to familiarise participants with the task. In the full experiment, each of the 13 orientations was presented 32 times in a randomised order, amounting to 416 trials in total. The experimental session lasted approximately 30 minutes.

### Quantifying the psychometric function

To model Gabor orientation discrimination performance, cumulative logistic psychometric functions (PFs) were fit to the data using a Maximum Likelihood criterion^82^. The dependent measure was the proportion of trials on which the participant indicated that the Gabor appeared to be oriented ‘rightward’ and the independent measure was the true orientation of the Gabor. The logistic function is described by the following:

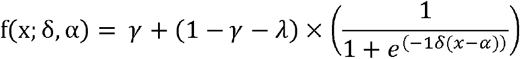

where ***x*** are the tested Gabor orientations, ***δ*** is the subjective threshold (location on the x-axis corresponding to 50% ‘left’/50% ‘right’ responses) and ***α*** is the slope of the rising curve (indexing visual sensitivity). Both ***λ*** and ***γ*** represent the probability of stimulus independent lapses and were fixed at 0.02.

### Data exclusion

The PF threshold and slope parameters were used to formally detect outliers in the dataset. Any participant who met any one of the following two criteria for the overall PF fit to their entire dataset was excluded from further analysis: 1) a threshold value over 3 median absolute deviations from the overall group median and/or 2) a slope value over 3 median absolute deviations from the overall group median. This led to a total of 6 participants being excluded and hence 37 participants were entered into the final inferential analyses.

### Quantifying perceptual choice history bias

In order to measure perceptual choice history bias, the data within each participant were split into two bins: one containing trials that followed a leftward orientation response on the previous trial (‘post left response’) and the other containing trials that followed a rightward orientation response on the previous trial (‘post right response’). The PF was fit separately to data from these subsets of trials (Fig. 2A). From the resulting fits, the threshold and slope were retrieved. This was done separately for trial lags of +1, +2 and +3. The difference in PF threshold between ‘post left’ and ‘post right’ responses indexes the strength and direction of perceptual choice history bias (Fig. 2B). If positive choice history bias (i.e. tendency to repeat previous choices^16–20^) heavily influences the orientation judgements then the group-averaged psychometric curves conditioned separately on ‘post left response’ and ‘post right response’ trials will be shifted horizontally on the *x*-axis in relation to one another. To formally test this, a repeated-measures *t*-test was performed to compare the PF thresholds between ‘post left response’ trials and ‘post right response’ trials. This analysis was also performed separately for ‘post high confidence’ and ‘post low confidence’ trials respectively (Supplementary Fig. 1: see *SDT parameter analyses* section below for division of confidence bins).

### Quantifying metacognitive choice history bias

Measuring history bias of metacognitive decisions required a different analytical approach. If positive metacognitive choice history bias occurs^31^ then confidence ratings will be more likely to be high following a high confidence rating and low following a low confidence rating, regardless of the level of external evidence (i.e. absolute Gabor orientation) (see Fig. 2C). To statistically test this, linear regression was performed between absolute Gabor orientation and mean confidence ratings separately for post 1, 2, 3 and 4 rating trials in each participant. Subsequently, linear regression was then performed between the previous confidence rating (1, 2, 3, 4) and the intercepts of the orientation-confidence regressions, and the resulting within-participant regression slope represented our measure of history bias. At the group level, a one sample *t*-test (versus 0) was performed on the resulting regression slopes to examine whether they were statistically different from zero (i.e. whether they showed a systematic directionality across participants). This was done separately for trial lags of +1, +2 and +3 (Fig. 2D). This analysis was also performed separately for trials in which the previous perceptual choice (i.e. ‘left’ or ‘right’) was ‘repeated’ versus trials in which the perceptual choice was ‘alternated’ (Fig 5A-B). A paired-samples t-test was used to compare the regression slopes between ‘repetition’ and ‘alternation’ trials.

### Quantifying choice history biases and estimating population prevalence using mutual information (MI)

Mutual Information (MI)^39^ is a measure of statistical dependence between two random variables that places no assumption on the form of the dependence. For two discrete variables *X* and *Y* that are distributed according to a joint probability distribution *P(X,Y)* the MI is defined as:

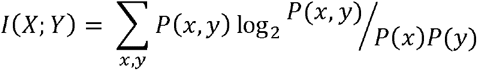

When the probability distributions are estimated from observed data the resulting MI estimate suffers from a limited sampling bias which causes the expectation of the estimate to be systematically larger than the true value. We correct this by subtracting the Miller-Madow bias estimate^83^, which is given by 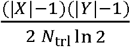, where |*X*|,|*Y*| are respectively the number of discrete values taken by the variables *X* and *Y*, and *N*_trl_ is the number of trials used for the calculation. Statistical inference was performed via permutation testing. The relationship between *X* and *Y* was shuffled and the resulting MI values stored. This was repeated 1000 times (separately for the each participant). The 95^th^ percentile of the resulting permutation value was used as the threshold for inference on the MI value obtained from the unshuffled data.

We calculate the following MI values (Figure 2E): I(Orient; Resp), I(Resp-1; Resp), I(Orient; Conf), I(Conf-1;Conf). In these calculations, the number of bins for the orientation is reduced by considering neighbouring levels of evidence together (e.g. 7 discrete bins corresponding to the following presented angles: [−18 −15] [−12 −9] [−6 −3] [0] [3 6] [9 12] [15 18]). Perceptual response is always represented with two discrete values (left or right). Confidence was represented with 3 or 4 discrete values (some participants never used one of the four confidence response values). For the choice history calculations, the variable Resp-1/Conf-1 is given by all trials excluding the last, the variable Resp/Conf is formed from all trials excluding the first.

### Modelling perceptual and metacognitive sensitivity and bias

Computational models of perceptual decision-making and confidence judgements, grounded largely in statistical decision theory and SDT, have successfully accounted for a range of confidence related empirical data^33,34,47,57^. Here, we modelled perceptual decisions and confidence ratings within an extended signal detection theory (SDT) framework^5^. This model assumes that, during yes/no detection or 2-alternative forced choice (2-AFC) discrimination tasks, binary decisions are made by the comparison of internal evidence (indexed by a noisy decision variable (***dv***)) with a decision criterion (***c***). Across trials, evidence generated by each stimulus class (i.e. noise/signal, choice A/choice B) is sampled from a stimulus-specific normal distribution. The relative separation between the distributions (in standard deviation units) indexes the overall level of evidence available for the decisions (***d’***), and hence how well the observer can discriminate between noise and signal, or between choice A and choice B. On a given trial, the probability that the choice is correct is indexed by the absolute distance between ***dv*** and ***c*** (in an unbiased observer), and hence statistically optimal confidence judgements should reflect this computation^34,47^. When a discrete confidence rating scale is employed, the rating on a given trial is defined by where the ***dv*** falls with respect to the so-called ‘type-2’ criteria (***c2***). The ***c2*** are response conditional, with separate criteria for the 2 possible choices (i.e. noise/signal, choice A/choice B). Overall, there are (k-1) × 2 ***c2***, where k equals the number of confidence ratings available. Figure 1B presents the model schematically for 3 differing levels of decision evidence: no evidence (left panel), weak evidence (middle panel) and strong evidence (right panel). The distributions and predicted effects in Figures 1B-E were produced using code developed by Urai et al.,^16^ (https://github.com/anne-urai/pupilUncertainty). The x-axis ranges from [−15:15] in these examples, ***d’*** was set to 0.1 (no evidence), 1.58 (weak evidence) and 3.17 (strong evidence) whereas c was always set to 0 (unbiased observer). The flanking ***c2*** were set at ±3 (conservative) and ±6 (liberal) for each. To formalise the predicted relationships between evidence strength, accuracy and confidence (Fig. 1-E), we simulated a normal distribution of ***dv*** for one response (i.e. *μ*>0) at each level of evidence strength. All samples from the simulated distribution were split into correct and error ‘choices’ based on their position relative to ***c***. For each combination of evidence strength and choice, the level of confidence is

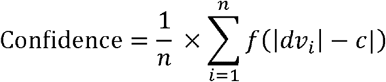

where *f* is the cumulative distribution function of the normal distribution

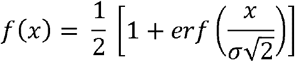

which transforms the distance between dv and c into the probability of a correct response^16,84^. Ten millions trials were simulated and for each iteration a binary choice was computed along with its accuracy and corresponding level of confidence. Because response times are often taken as a proxy of decision confidence (with response times increasing as a function of decreasing confidence)^16,34^ the response time prediction (Fig. 1E) represents an inversion of the confidence prediction (Fig. 1D).

In order to quantify both type-1 and type-2 performance parameters (i.e. sensitivity and bias) across different levels of evidence strength (absolute Gabor orientations) in the real data, we adopted the meta-d’ approach (see ^5,42,43^ for extended description and discussion) as implemented using single-subject Bayesian model fits within the ‘HMeta-d’ toolbox^42^ (https://github.com/metacoglab/HMeta-d). ***Meta-d’*** characterises type-2 sensitivity as the value of ***d’*** that a metacognitively optimal observer, with the same type-1 response bias (**c**), would have required to produce the observed type-2 (confidence) data^5^. If an observer has perfect metacognitive insight (i.e. they are always high in confidence when correct and low in confidence when incorrect) then ***d’*** will be equal to ***meta-d’***. Importantly, because ***meta-d’*** is expressed in the same units as ***d’***, the two can be compared directly to quantify the level of metacognitive efficiency/insight. If the metacognitive efficiency score (***meta- d’-d’***) ≠ 0, then the type-2 responses (confidence ratings) are either more (positive value) or less (negative value) sensitive to the task-related evidence than the type-1 perceptual responses. We note that (***meta-d’/d’***) is often used to quantify metacognitive efficiency as a ratio of type-1 performance^58^ and so we replicated our correlation analyses involving (***meta-d’-d’***) using (***meta- d’/d’***) (see Supplementary Figure 4). The same pattern of results was found. The metacognitive criteria (***meta-c***) represent type-2 bias (***c2***) calculated within the meta-d’ framework: the tendency to give high or low confidence ratings regardless of evidence strength. We calculated the absolute distance between ***meta-c*** and type-1 ***c*** (**|*meta-c - c*|**) in order to isolate the metacognitive response bias from the perceptual response bias^43^. Lower values of **|*meta-c - c*|** indicate an overall response bias in favour of higher confidence ratings. As mentioned, ***meta-c (c2)*** values are calculated separately for each of the possible perceptual responses (i.e. ‘left’ or ‘right’ orientation judgements in the current study) and for each of N-1 confidence ratings available to choose from (4 in the current experiment). In order to streamline the analysis, we averaged over the 3 **|*meta-c - c*|** values for each response (‘left’ or ‘right’) separately to gain a single estimate of overall metacognitive response bias.

### Statistical analyses on SDT parameters

We compared overall perceptual sensitivity (***d’***) to metacognitive sensitivity (***meta-d’***) across all levels of evidence strength using a 2 (sensitivity measure: ***d’, meta-d’***) × 6 (absolute Gabor orientation: 3°,6°,9°,12°,15°,18°) repeated measures ANOVA. To assess the extent to which the type-1 and type-2 SDT performance parameters were influenced by both perceptual and metacognitive choice history, trials were binned in three different ways (1. ‘post left’/’post right’ choice trials (Fig. 3); 2. ‘post high’/’post low’ confidence trials (Fig. 4); 3. ‘repetition’/’alternation’ trials (Fig. 5)) and the parameters (***d’, meta-d’, meta-d’ – d’, c, |meta-c - c|:‘left’ responses, |meta-c - c|:’right’ responses***) were calculated for both bins separately at each of the 6 levels of evidence strength. Repeated measures ANOVAs (2 (choice history bin) × 6 (absolute Gabor orientation: 3°,6°,9°,12°,15°,18°)) were then performed separately for each parameter. Significant interaction terms were followed up using paired samples t-tests of the difference between the choice history bins separately at each level of evidence strength. To split the trials into relatively equal ‘post high’ and ‘post low’ confidence bins within each participant, the number of trials immediately following each of the 4 confidence ratings (i.e. post ‘1’, ‘2’, ‘3’, ‘4’ ratings) was calculated and bins were assigned that minimized the difference in trial number between the high and low bins (median difference between bins = 69 trials (min = 7, max = 251)). This led to 10 participants having ‘low’ bin = ‘1’, ‘2’ and ‘3’ ratings, ‘high’ bin = ‘4’ ratings, 14 participants (‘low’ bin = ‘1’ and ‘2’ ratings, ‘high’ bin = ‘3’ and ‘4’ ratings) and 13 participants (‘low’ bin = ‘1’ ratings, ‘high’ bin = ‘2’, ‘3’ and ‘4’ ratings). Note that 4 participants were excluded from the analysis of the influence of previous confidence level on perceptual choice history bias (Supplementary Fig. 1) because they had PF slope values over 3 median absolute deviations from the overall group median in at least one of the conditions here. This was due to biased perceptual and/or confidence decisions leading to a small number of trials being available for PF fitting after binning for these participants.

For all t-tests and correlations (see below), we calculated the Bayes Factor (BF_10_) obtained from paired-samples Bayesian t-tests^85^ or correlation hypothesis tests^86^, with a prior following a Cauchy distribution and a scale factor of 0.707. BF_10_ quantifies the evidence in favour of the null or alternative hypotheses, where BF_10_ below 1/3 indicates evidence for the null hypothesis, above 3 indicates evidence for the alternative hypothesis and between 1/3 and 3 indicates that the evidence is inconclusive (potentially due to a lack of statistical power)^85^.

## Between-subject correlations

Both Pearson and Spearman correlation coefficients were calculated for each of the between-subject correlations of interest. Only Pearson’s r are shown in the corresponding figures.

## Data Availability

All data and code to reproduce the analyses are openly available on the Open Science Framework (OSF) under the URL: https://osf.io/5chwq/

## Acknowledgements

CSYB was supported by the British Academy/Leverhulme Trust [SRG19/191169]. RAAI was supported by the Wellcome Trust [214120/Z/18/Z].

## Author Contributions

Conceptualization, C.S.Y.B.; Data collection, R.B. and F.W.; Formal Analysis, C.S.Y.B and R.A.A.I.; Writing, C.S.Y.B., R.B. and R.A.A.I. Supervision, C.S.Y.B.

## Supplementary Information

### Supplementary Figures

**Supplementary Figure 1.**
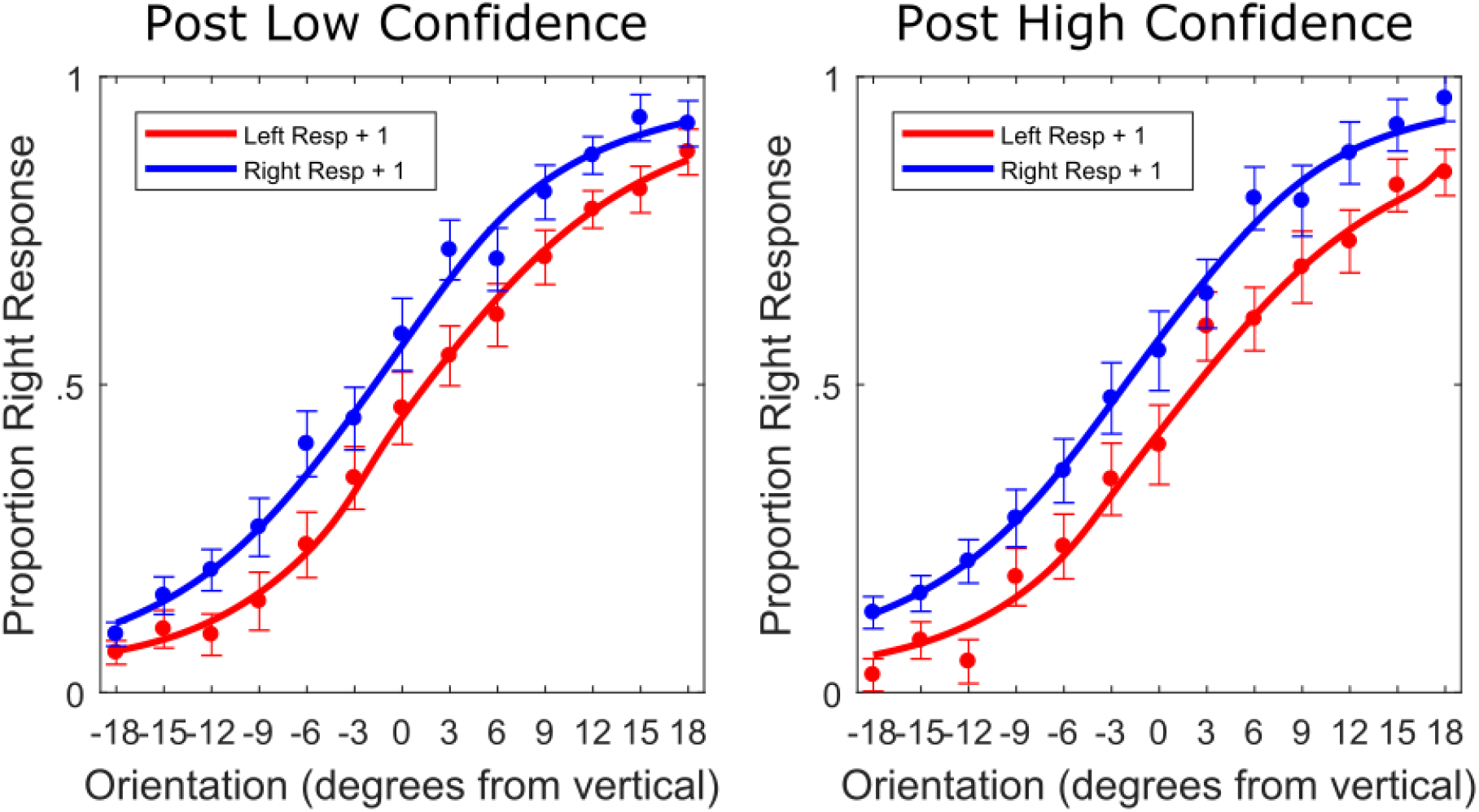
Prior confidence did not significantly alter the strength of perceptual choice history bias. The same figure as Fig. 2A but separately for ‘post low confidence’ and ‘post high confidence’ trials. Trials were split into post high and low confidence bins as evenly as possible within each participant (see Methods). Four participants had to be excluded because they did not have enough trials in one of the conditions to reliably retrieve psychometric function (PF) parameter estimates (see Methods). A 2 (previous confidence level: low versus high) × 2 (previous perceptual choice: left versus right) repeated-measures ANOVA on PF thresholds revealed a significant main effect of previous perceptual choice (F(1,32) = 11.584, p = .002, 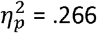), no main effect of previous confidence level (F(1,32) = .891, p = .352, 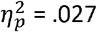) and no interaction between the two (F(1,32) = 1.689, p = .203, 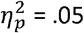). Post ‘left choice’ thresholds were significantly different to post ‘right choice’ thresholds, both for post low confidence trials (t(32) = 4.029, p < .001, BF_10_ = 88.338) and for post high confidence trials (t(32) = 2.781, p = .009, BF_10_ = 4.765). The difference in perceptual choice history bias (post ‘left choice’ – post ‘right choice’ thresholds) between post high and post low confidence trials was not significant (t(32) = 1.3, p = .203, BF_10_ = 0.402). These results are in apparent contrast to those of previous studies^1–4^. For instance, Braun et al^2^ showed that confidence (indexed by RT and accuracy) boosted adaptation to experimentally controlled changes in the repetition probability of the stimuli. Here we did not manipulate repetition probability which remained random throughout. We also employed explicit reports of confidence whereas previous studies have employed RT, accuracy and/or pupil diameter as proxies for confidence/uncertainty^1,2^. In another study involving explicit reports, Samaha et al^4^ employed a task which dissociated confidence ratings from objective performance and found that when sensory evidence in favour of one choice versus another was kept constant, higher confidence trials led to stronger serial dependence than lower confidence trials. Hence, ‘pure’ fluctuations of confidence occurring irrespective of relative choice evidence may influence history bias^4^, whereas the effect may be ameliorated when most ratings are informed by the quality of evidence (as was the case here). In other words, the interplay between perceptual decisions, confidence and choice history bias might vary depending on the information available to inform decisions and confidence judgements. This hypothesis could be directly tested by manipulating the quality of evidence available for perceptual choices whilst matching task accuracy^4,5^ within different blocks of the same experiment. It would also be instructive for future studies to tease apart and compare the relative contributions of different measures which putatively index uncertainty/confidence (such as explicit ratings, RT, event-related potentials^6,7^ and pupil diameter) but which may exert dissociable influences on perceptual and metacognitive history biases^1^.

**Supplementary Figure 2.**
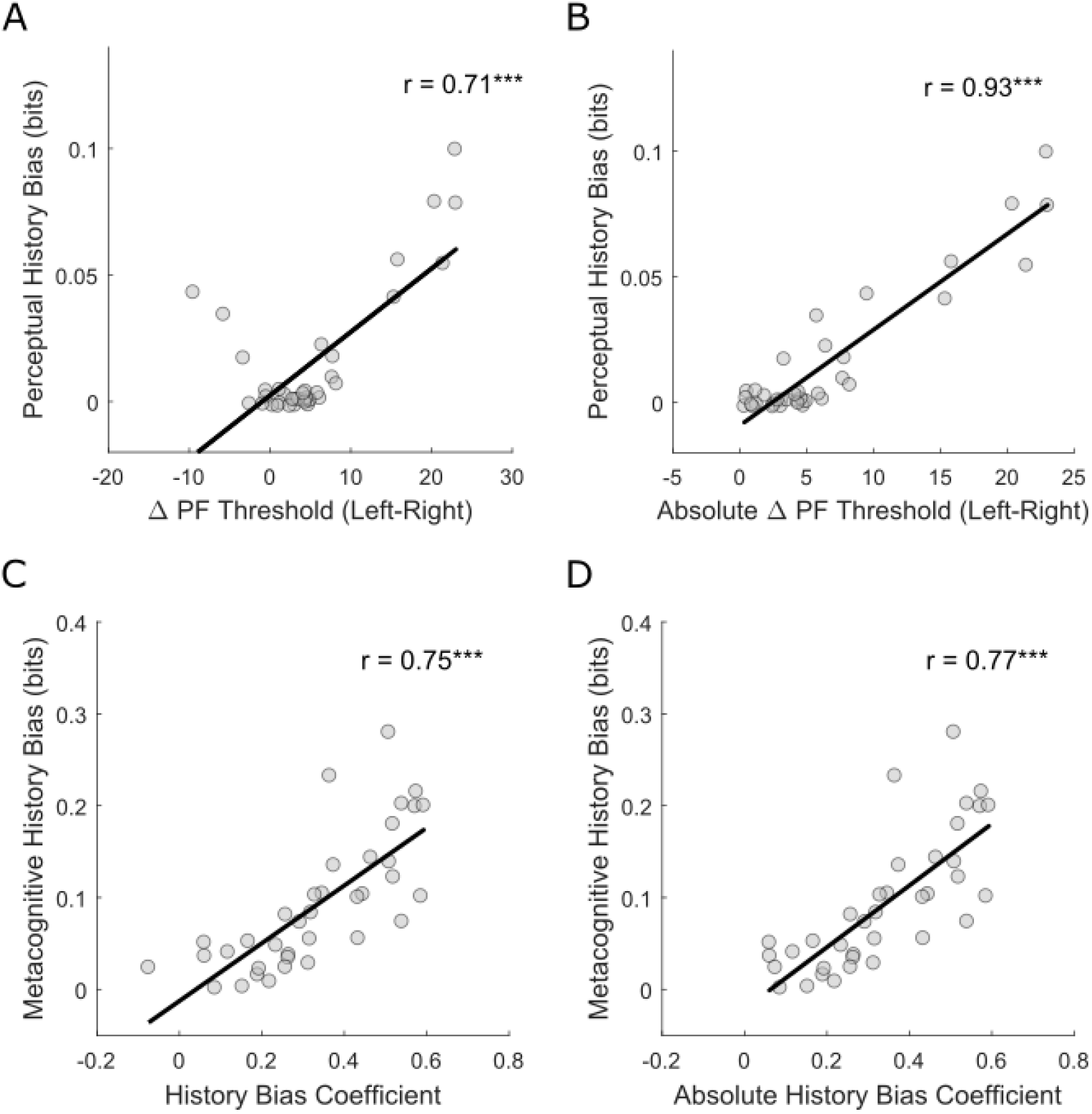
Correlations between model-based and non-parametric measures of choice history bias. The MI history bias measure is an assumption free measure which indexes how strongly current responses are related to previous responses, irrespective of the evidence available on a given trial. **(a)** Relationship between differences in psychometric function (PF) threshold between ‘post-left choice’ and ‘post-right choice’ trials and perceptual history bias quantified using mutual information (I(resp-1;resp)) (Pearson’s r = 0.7124, p < .001, Spearman’s rho = 0.4697, p = .004, BF_10_ = 2.3173e+04). Notice the expected u-shaped relationship which occurs because MI is an unsigned quantification of dependence (i.e. it does not dissociate ‘repetition’ from ‘alternation’ biases). **(b)** Relationship between absolute differences in psychometric function (PF) threshold between ‘post-left choice’ and ‘post-right choice’ trials and perceptual history bias quantified using mutual information (I(resp-1;resp)) (Pearson’s r = 0.9292, p < .001, Spearman’s rho = 0.725, p < .001, BF_10_ = 6.9779e+13). The absolute strength of the PF difference bias (regardless of sign) is strongly linearly related to the MI measure of bias. **(c)** Relationship between metacognitive history bias coefficients and metacognitive history bias quantified using mutual information (I(conf-1;conf)) (Pearson’s r = 0.7516, p < .001, Spearman’s rho = 0.8286, p < .001, BF_10_ = 1.9176e+05). **(d)** Relationship between absolute metacognitive history bias coefficients and metacognitive history bias quantified using mutual information (I(conf-1;conf)) (Pearson’s r = 0.7654, p < .001, Spearman’s rho = 0.8257, p < .001, BF_10_ = 4.4338e+05).

**Supplementary Figure 3.**
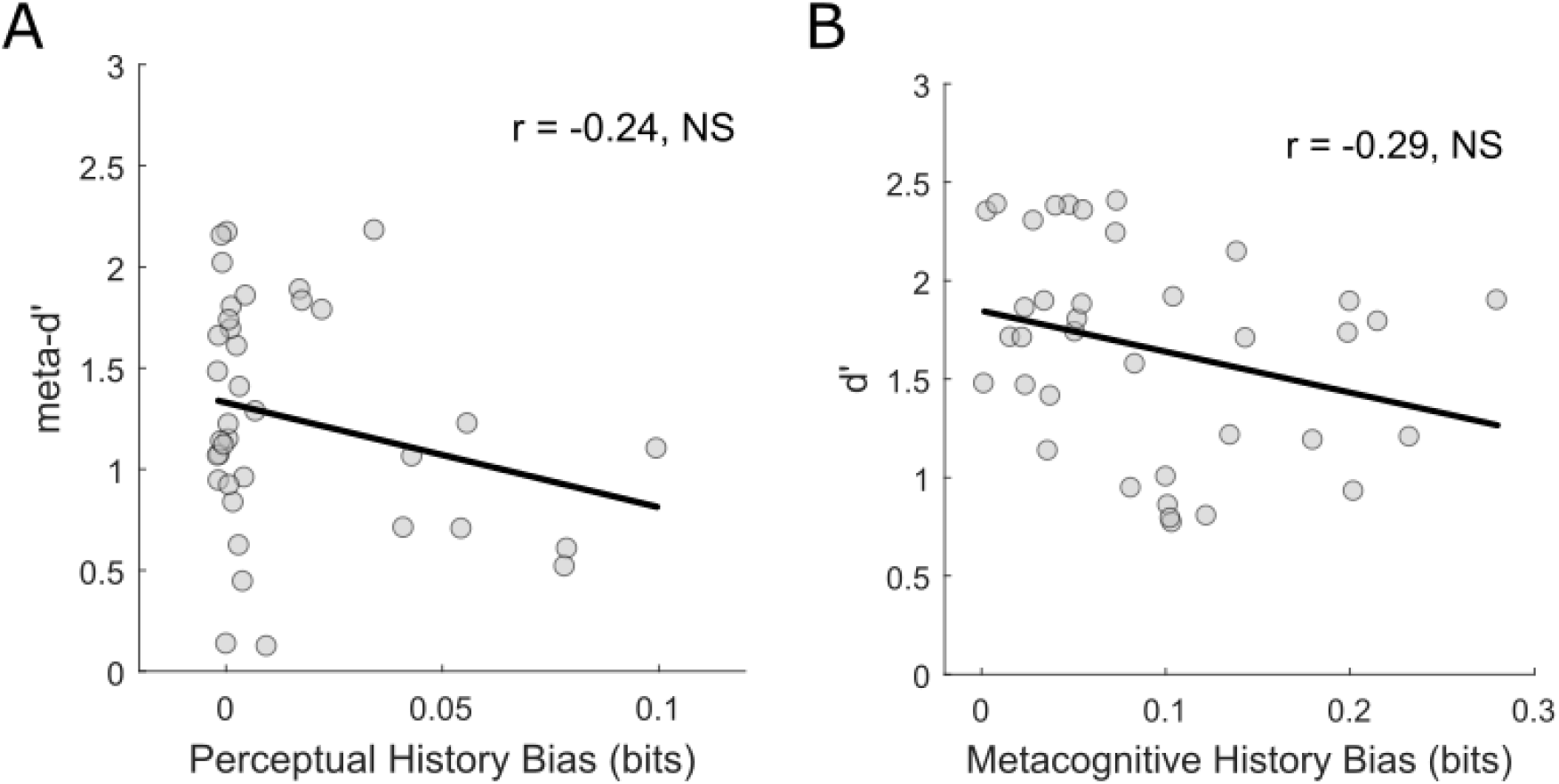
Between-subjects Pearson correlations. **(a)** Relationship between perceptual choice history bias and metacognitive sensitivity (***meta-d’***) (Pearson’s r = −0.245, p = .1439, Spearman’s rho = −0.1802, p = .2847, BF_10_ = 0.369). **(b)** Relationship between metacognitive choice history bias and perceptual sensitivity (***d’***) (Pearson’s r = −0.2874, p = .0846, Spearman’s rho = −0.2992, p = .0724, BF_10_ = 0.56).

**Supplementary Figure 4.**
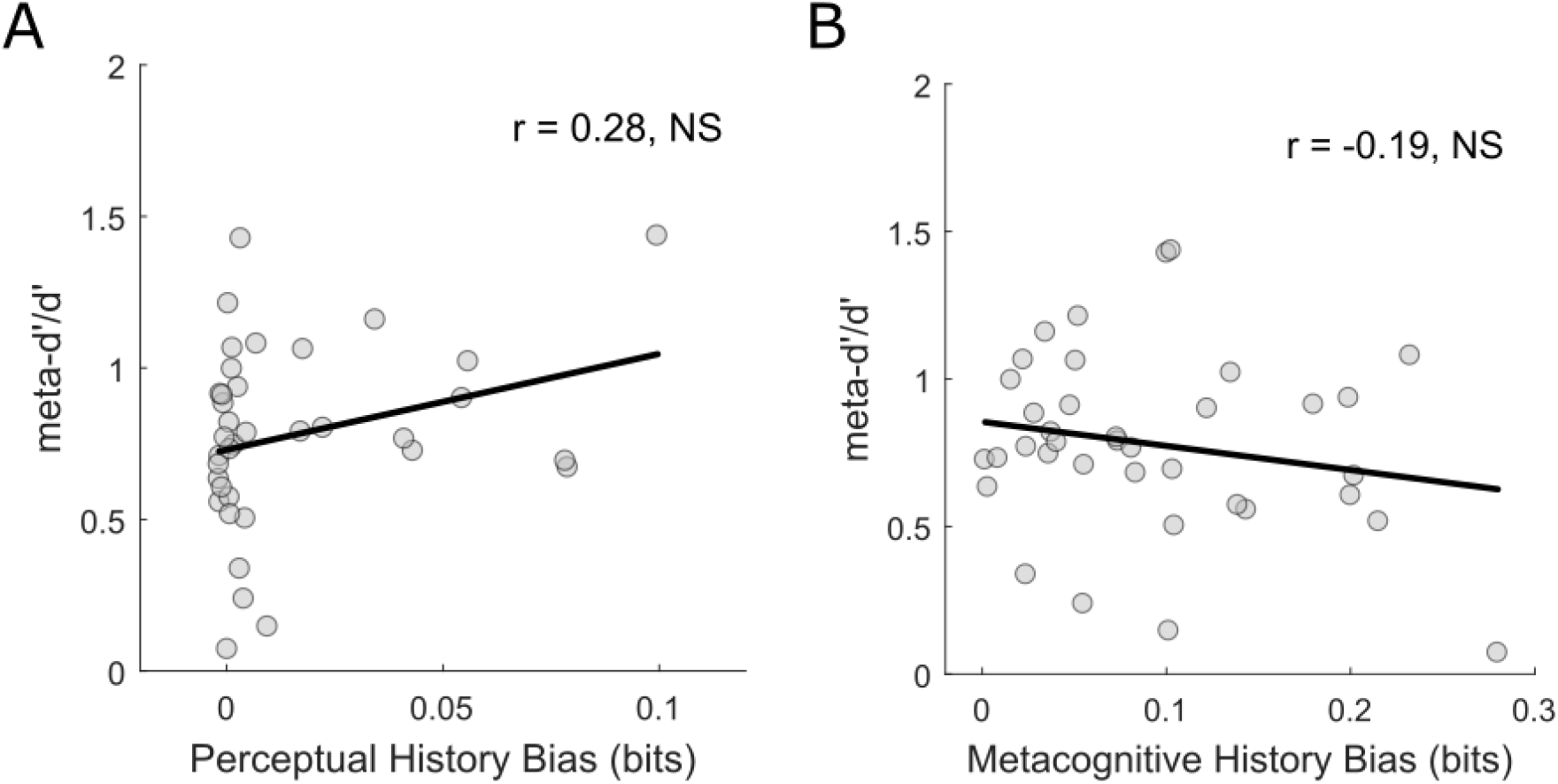
Between-subjects metacognitive insight ratio correlations. **(a)** Relationship between perceptual choice history bias and metacognitive efficiency (***meta-d’/d’***) (Pearson’s r = 0.2759, p = .0983, Spearman’s rho = 0.2349, p = .1612, BF_10_ = 0.497). **(b)** Relationship between metacognitive choice history bias and metacognitive efficiency (***meta-d’/d’***) (Pearson’s r = −0.1944, p = .249, Spearman’s rho = −0.169, p = .316, BF_10_ = 0.247).

**Supplementary Figure 5.**
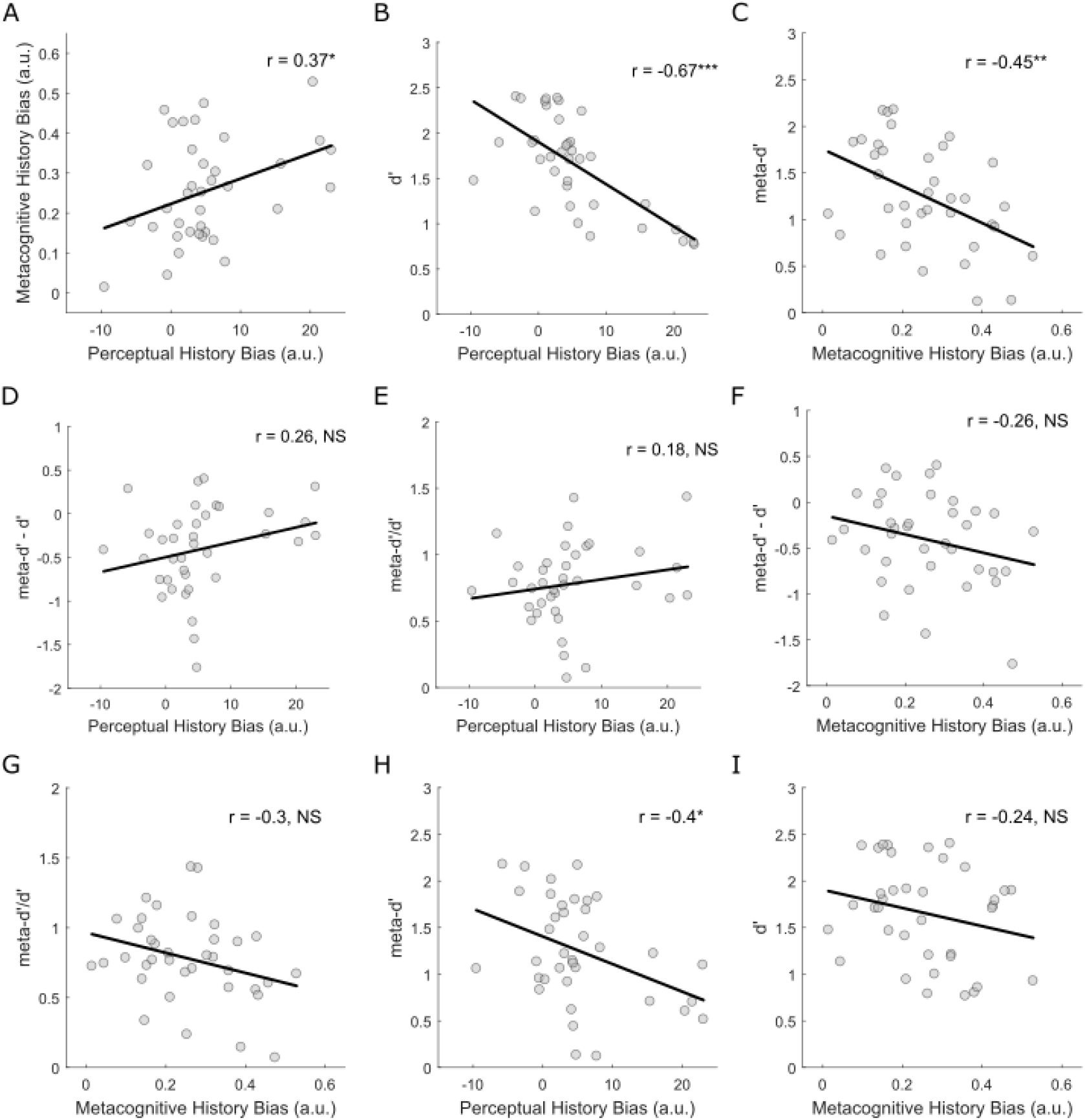
Between-subjects Pearson correlations using metrics of choice history bias which retain the direction of bias (i.e. repeater (+ve x-axis values) versus alternator (−ve x-axis values)). Perceptual choice history bias was quantified for each participant by subtracting their post left choice PF threshold from their post right choice PF threshold. Metacognitive choice history bias was quantified by a linear regression coefficient linking confidence ratings on trials 1:415 and confidence ratings on trials 2:416. **(a)** Relationship between perceptual and metacognitive choice history biases (Pearson’s r = 0.3723, p = .0232, Spearman’s rho = 0.2667, p = .1105, BF_10_ = 1.647). The positive correlation suggests that the stronger the positive perceptual history bias (i.e. ‘repeaters’), the stronger the metacognitive choice history bias. However, though the Pearson correlation was significant, Spearman’s Rho was not significant and BF_10_ indicated that the data were insensitive as to whether the effect exists. Hence, this effect should be treated with caution and we do not interpret it further here. **(b)** Relationship between perceptual choice history bias and perceptual sensitivity (***d’***) (Pearson’s r = −0.6725, p < .001, Spearman’s rho = −0.6385, p < .001, BF_10_ = 3.7753e+03). Stronger perceptual repetition biases were associated with reduced perceptual sensitivity. **(c)** Relationship between metacognitive choice history bias and metacognitive sensitivity (***meta-d’***) (Pearson’s r = −0.4525, p = .005, Spearman’s rho = −0.426, p = .009, BF_10_ = 6.462). Stronger metacognitive repetition biases were associated with reduced metacognitive sensitivity. **(d)** Relationship between perceptual choice history bias and metacognitive efficiency (indexed by ***meta-d’ – d’***) (Pearson’s r = 0.26, p = .12, Spearman’s rho = −0.3872, p = .018, BF_10_ = 0.425). The positive correlation suggests that the stronger the positive perceptual history bias (i.e. ‘repeaters’), the higher the level of metacognitive insight. However, though the Spearman correlation was significant, Pearson’s *r* was not significant and BF_10_ indicated that the data were insensitive as to whether the effect exists. **(e)** Relationship between perceptual choice history bias and metacognitive efficiency (indexed by ***meta-d’/d’***) (Pearson’s r = 0.18, p = .285, Spearman’s rho = 0.2378, p = .156, BF_10_ = 0.2255). **(f)** Relationship between metacognitive choice history bias and metacognitive efficiency (indexed by ***meta-c – d’***) (Pearson’s r = −0.26, p = .1198, Spearman’s rho = −0.2278, p = .1745, BF_10_ = 0.4252). **(g)** Relationship between metacognitive choice history bias and metacognitive efficiency (indexed by ***meta-d’/d’***) (Pearson’s r = −0.3031, p = .0682, Spearman’s rho = −0.3025, p = .0691, BF_10_ = 0.6662). **(h)** Relationship between perceptual choice history bias and metacognitive sensitivity (***meta-d’***) (Pearson’s r = −0.3978, p = .0148, Spearman’s rho = −0.3227, p = .0519, BF_10_ = 2.4422). The negative correlation suggests that the stronger the perceptual repetition bias, the lower the metacognitive sensitivity. However, though the Pearson correlation was significant, Spearman’s Rho was not significant and BF_10_indicated that the data were insensitive as to whether the effect exists. **(i)** Relationship between metacognitive choice history bias and perceptual sensitivity (***d’***) (Pearson’s r = −0.2388, p = .1546, Spearman’s rho = −0.2148, p = .2010, BF_10_ = 0.3498).

## References

1 Wilson, T. D. & Dunn, E. W. Self-knowledge: its limits, value, and potential for improvement. Annual review of psychology 55, 493–518, doi:10.1146/annurev.psych.55.090902.141954 (2004).

2 Johansson, P., Hall, L., Sikstrom, S. & Olsson, A. Failure to detect mismatches between intention and outcome in a simple decision task. Science 310, 116–119, doi:10.1126/science.1111709 (2005).

3 Johnson, D. D. & Fowler, J. H. The evolution of overconfidence. Nature 477, 317–320, doi:10.1038/nature10384 (2011).

4 Galvin, S. J., Podd, J. V., Drga, V. & Whitmore, J. Type 2 tasks in the theory of signal detectability: discrimination between correct and incorrect decisions. Psychonomic bulletin & review 10, 843–876 (2003).

5 Maniscalco, B. & Lau, H. A signal detection theoretic approach for estimating metacognitive sensitivity from confidence ratings. Conscious Cogn 21, 422–430, doi:10.1016/j.concog.2011.09.021 (2012).

6 Fleming, S. M. & Lau, H. C. How to measure metacognition. Frontiers in human neuroscience 8, 443, doi:10.3389/fnhum.2014.00443 (2014).

7 Fleming, S. M., Weil, R. S., Nagy, Z., Dolan, R. J. & Rees, G. Relating introspective accuracy to individual differences in brain structure. Science 329, 1541–1543, doi:10.1126/science.1191883 (2010).

8 Peters, M. A. K. et al. Perceptual confidence neglects decision-incongruent evidence in the brain. Nat Hum Behav 1, doi:10.1038/s41562-017-0139 (2017).

9 van den Berg, R., Zylberberg, A., Kiani, R., Shadlen, M. N. & Wolpert, D. M. Confidence Is the Bridge between Multi-stage Decisions. Current biology: CB 26, 3157–3168, doi:10.1016/j.cub.2016.10.021 (2016).

10 Desender, K., Boldt, A. & Yeung, N. Subjective Confidence Predicts Information Seeking in Decision Making. Psychological science 29, 761–778, doi:10.1177/0956797617744771 (2018).

11 Yeung, N. & Summerfield, C. Metacognition in human decision-making: confidence and error monitoring. Philosophical transactions of the Royal Society of London. Series B, Biological sciences 367, 1310–1321, doi:10.1098/rstb.2011.0416 (2012).

12 Bahrami, B. et al. What failure in collective decision-making tells us about metacognition. Philosophical transactions of the Royal Society of London. Series B, Biological sciences 367, 1350–1365, doi:10.1098/rstb.2011.0420 (2012).

13 Folke, T., Jacobsen, C., Fleming, S. M. & De Martino, B. Explicit representation of confidence informs future value-based decisions. Nature Human Behaviour 1 (2016).

14 David, A. S., Bedford, N., Wiffen, B. & Gilleen, J. Failures of metacognition and lack of insight in neuropsychiatric disorders. Philosophical transactions of the Royal Society of London. Series B, Biological sciences 367, 1379–1390, doi:10.1098/rstb.2012.0002 (2012).

15 Rouault, M., Seow, T., Gillan, C. M. & Fleming, S. M. Psychiatric Symptom Dimensions Are Associated With Dissociable Shifts in Metacognition but Not Task Performance. Biol Psychiatry 84, 443–451, doi:10.1016/j.biopsych.2017.12.017 (2018).

16 Urai, A. E., Braun, A. & Donner, T. H. Pupil-linked arousal is driven by decision uncertainty and alters serial choice bias. Nature communications 8, 14637, doi:10.1038/ncomms14637 (2017).

17 Braun, A., Urai, A. E. & Donner, T. H. Adaptive History Biases Result from Confidence-weighted Accumulation of Past Choices. The Journal of neuroscience: the official journal of the Society for Neuroscience, doi:10.1523/JNEUROSCI.2189-17.2017 (2018).

18 Bonaiuto, J. J., Berker, A. & Bestmann, S. Response repetition biases in human perceptual decisions are explained by activity decay in competitive attractor models. eLife 5, doi:10.7554/eLife.20047 (2016).

19 Abrahamyan, A., Silva, L. L., Dakin, S. C., Carandini, M. & Gardner, J. L. Adaptable history biases in human perceptual decisions. Proceedings of the National Academy of Sciences of the United States of America 113, E3548–3557, doi:10.1073/pnas.1518786113 (2016).

20 Urai, A. E., de Gee, J. W., Tsetsos, K. & Donner, T. H. Choice history biases subsequent evidence accumulation. eLife 8, doi:10.7554/eLife.46331 (2019).

21 Fernberger, S. W. Interdependence of judgments within the series for the method of constant stimuli. J Exp Psychol 3, 126–150, doi:DOI 10.1037/h0065212 (1920).

22 Fritsche, M., Mostert, P. & de Lange, F. P. Opposite Effects of Recent History on Perception and Decision. Current biology: CB 27, 590–595, doi:10.1016/j.cub.2017.01.006 (2017).

23 Fischer, J. & Whitney, D. Serial dependence in visual perception. Nature neuroscience 17, 738–743, doi:10.1038/nn.3689 (2014).

24 Bliss, D. P., Sun, J. J. & D’Esposito, M. Serial dependence is absent at the time of perception but increases in visual working memory. Scientific reports 7, 14739, doi:10.1038/s41598-017-15199-7 (2017).

25 Liberman, A., Fischer, J. & Whitney, D. Serial dependence in the perception of faces. Current biology: CB 24, 2569–2574, doi:10.1016/j.cub.2014.09.025 (2014).

26 St John-Saaltink, E., Kok, P., Lau, H. C. & de Lange, F. P. Serial Dependence in Perceptual Decisions Is Reflected in Activity Patterns in Primary Visual Cortex. J Neurosci 36, 6186–6192, doi:10.1523/JNEUROSCI.4390-15.2016 (2016).

27 Pascucci, D. et al. Laws of concatenated perception: Vision goes for novelty, decisions for perseverance. PLoS biology 17, doi:ARTN e3000144 10.1371/journal.pbio.3000144 (2019).

28 Kiyonaga, A., Scimeca, J. M., Bliss, D. P. & Whitney, D. Serial Dependence across Perception, Attention, and Memory. Trends Cogn Sci 21, 493–497, doi:10.1016/j.tics.2017.04.011 (2017).

29 Cicchini, G. M., Mikellidou, K. & Burr, D. C. The functional role of serial dependence. Proceedings. Biological sciences 285, doi:10.1098/rspb.2018.1722 (2018).

30 Manassi, M., Liberman, A., Chaney, W. & Whitney, D. The perceived stability of scenes: serial dependence in ensemble representations. Scientific reports 7, 1971, doi:10.1038/s41598-017-02201-5 (2017).

31 Rahnev, D., Koizumi, A., McCurdy, L. Y., D’Esposito, M. & Lau, H. Confidence Leak in Perceptual Decision Making. Psychological science 26, 1664–1680, doi:10.1177/0956797615595037 (2015).

32 Samaha, J., Switzky, M. & Postle, B. R. Confidence boosts serial dependence in orientation estimation. J Vision 19, doi:Artn 25 10.1167/19.4.25 (2019).

33 Kepecs, A., Uchida, N., Zariwala, H. A. & Mainen, Z. F. Neural correlates, computation and behavioural impact of decision confidence. Nature 455, 227–231, doi:10.1038/nature07200 (2008).

34 Sanders, J. I., Hangya, B. & Kepecs, A. Signatures of a Statistical Computation in the Human Sense of Confidence. Neuron 90, 499–506, doi:10.1016/j.neuron.2016.03.025 (2016).

35 Hebart, M. N., Schriever, Y., Donner, T. H. & Haynes, J. D. The Relationship between Perceptual Decision Variables and Confidence in the Human Brain. Cerebral cortex 26, 118–130, doi:10.1093/cercor/bhu181 (2016).

36 Hangya, B., Sanders, J. I. & Kepecs, A. A Mathematical Framework for Statistical Decision Confidence. Neural Comput 28, 1840–1858, doi:10.1162/NECO_a_00864 (2016).

37 Drugowitsch, J. Becoming Confident in the Statistical Nature of Human Confidence Judgments. Neuron 90, 425–427, doi:10.1016/j.neuron.2016.04.023 (2016).

38 Fleming, S. M. & Daw, N. D. Self-Evaluation of Decision-Making: A General Bayesian Framework for Metacognitive Computation. Psychol Rev 124, 91–114, doi:10.1037/rev0000045 (2017).

39 Ince, R. A. et al. A statistical framework for neuroimaging data analysis based on mutual information estimated via a gaussian copula. Hum Brain Mapp 38, 1541–1573, doi:10.1002/hbm.23471 (2017).

40 Donhauser, P. W., Florin, E. & Baillet, S. Imaging of neural oscillations with embedded inferential and group prevalence statistics. PLoS computational biology 14, doi:ARTN e1005990 10.1371/journal.pcbi.1005990 (2018).

41 Allefeld, C., Gorgen, K. & Haynes, J. D. Valid population inference for information-based imaging: From the second-level t-test to prevalence inference. NeuroImage 141, 378–392, doi:10.1016/j.neuroimage.2016.07.040 (2016).

42 Fleming, S. M. HMeta-d: hierarchical Bayesian estimation of metacognitive efficiency from confidence ratings. Neuroscience of consciousness 2017, nix007, doi:10.1093/nc/nix007 (2017).

43 Sherman, M. T., Seth, A. K. & Barrett, A. B. Quantifying metacognitive thresholds using signal-detection theory. bioRxiv (2018).

44 Drugowitsch, J., Moreno-Bote, R. & Pouget, A. Relation between Belief and Performance in Perceptual Decision Making. PloS one 9, doi:ARTN e96511 10.1371/journal.pone.0096511 (2014).

45 Rahnev, D. & Denison, R. N. Suboptimality in Perceptual Decision Making. The Behavioral and brain sciences, 1–107, doi:10.1017/S0140525X18000936 (2018).

46 Wyart, V. & Koechlin, E. Choice variability and suboptimality in uncertain environments. Curr Opin Behav Sci 11, 109–115, doi:10.1016/j.cobeha.2016.07.003 (2016).

47 Pouget, A., Drugowitsch, J. & Kepecs, A. Confidence and certainty: distinct probabilistic quantities for different goals. Nature neuroscience 19, 366–374, doi:10.1038/nn.4240 (2016).

48 Gherman, S. & Philiastides, M. G. Neural representations of confidence emerge from the process of decision formation during perceptual choices. NeuroImage 106, 134–143, doi:10.1016/j.neuroimage.2014.11.036 (2015).

49 Kiani, R. & Shadlen, M. N. Representation of Confidence Associated with a Decision by Neurons in the Parietal Cortex. Science 324, 759–764, doi:10.1126/science.1169405 (2009).

50 Meyniel, F., Schlunegger, D. & Dehaene, S. The Sense of Confidence during Probabilistic Learning: A Normative Account. PLoS computational biology 11, doi:ARTN e1004305 10.1371/journal.pcbi.1004305 (2015).

51 van den Berg, R. et al. A common mechanism underlies changes of mind about decisions and confidence. eLife 5, e12192, doi:10.7554/eLife.12192 (2016).

52 Huettel, S. A., Song, A. W. & McCarthy, G. Decisions under uncertainty: Probabilistic context influences activation of prefrontal and parietal cortices. Journal of Neuroscience 25, 3304–3311, doi:DOI 10.1523/Jneurosci.5070-04.2005 (2005).

53 Bang, D. et al. Confidence matching in group decision-making. Nature Human Behaviour 1, doi:UNSP 0117 10.1038/s41562-017-0117 (2017).

54 Denison, R. N., Adler, W. T., Carrasco, M. & Ma, W. J. Humans incorporate attention-dependent uncertainty into perceptual decisions and confidence. Proceedings of the National Academy of Sciences of the United States of America 115, 11090–11095, doi:10.1073/pnas.1717720115 (2018).

55 Rahnev, D. et al. Attention induces conservative subjective biases in visual perception. Nature neuroscience 14, 1513–1515, doi:10.1038/nn.2948 (2011).

56 Maniscalco, B., McCurdy, L. Y., Odegaard, B. & Lau, H. Limited Cognitive Resources Explain a Trade-Off between Perceptual and Metacognitive Vigilance. Journal of Neuroscience 37, 1213–1224, doi:10.1523/Jneurosci.2271-13.2016 (2017).

57 Pleskac, T. J. & Busemeyer, J. R. Two-Stage Dynamic Signal Detection: A Theory of Choice, Decision Time, and Confidence. Psychol Rev 117, 864–901, doi:10.1037/a0019737 (2010).

58 Morales, J., Lau, H. & Fleming, S. M. Domain-General and Domain-Specific Patterns of Activity Supporting Metacognition in Human Prefrontal Cortex. Journal of Neuroscience 38, 3534–3546, doi:10.1523/Jneurosci.2360-17.2018 (2018).

59 Fleming, S. M., Huijgen, J. & Dolan, R. J. Prefrontal Contributions to Metacognition in Perceptual Decision Making. Journal of Neuroscience 32, 6117–6125, doi:10.1523/Jneurosci.6489-11.2012 (2012).

60 Bang, D. & Fleming, S. M. Distinct encoding of decision confidence in human medial prefrontal cortex. Proceedings of the National Academy of Sciences of the United States of America 115, 6082–6087, doi:10.1073/pnas.1800795115 (2018).

61 Lebreton, M., Abitbol, R., Daunizeau, J. & Pessiglione, M. Automatic integration of confidence in the brain valuation signal. Nature neuroscience 18, 1159–+, doi:10.1038/nn.4064 (2015).

62 Murphy, P. R., Robertson, I. H., Harty, S. & O’Connell, R. G. Neural evidence accumulation persists after choice to inform metacognitive judgments. eLife 4, doi:ARTN e11946 10.7554/eLife.11946 (2015).

63 De Martino, B., Fleming, S. M., Garrett, N. & Dolan, R. J. Confidence in value-based choice. Nature neuroscience 16, 105–U147, doi:10.1038/nn.3279 (2013).

64 Benwell, C. S. Y. et al. Prestimulus EEG Power Predicts Conscious Awareness But Not Objective Visual Performance. eneuro 4, ENEURO.0182-0117.2017, doi:10.1523/eneuro.0182-17.2017 (2017).

65 Zylberberg, A., Roelfsema, P. R. & Sigman, M. Variance misperception explains illusions of confidence in simple perceptual decisions. Consciousness and Cognition 27, 246–253, doi:10.1016/j.concog.2014.05.012 (2014).

66 Ais, J., Zylberberg, A., Barttfeld, P. & Sigman, M. Individual consistency in the accuracy and distribution of confidence judgments. Cognition 146, 377–386, doi:10.1016/j.cognition.2015.10.006 (2016).

67 Rounis, E., Maniscalco, B., Rothwell, J. C., Passingham, R. E. & Lau, H. Theta-burst transcranial magnetic stimulation to the prefrontal cortex impairs metacognitive visual awareness. Cogn Neurosci-Uk 1, 165–175, doi:10.1080/17588921003632529 (2010).

68 Rahnev, D., Maniscalco, B., Luber, B., Lau, H. & Lisanby, S. H. Direct injection of noise to the visual cortex decreases accuracy but increases decision confidence. Journal of neurophysiology 107, 1556–1563, doi:10.1152/jn.00985.2011 (2012).

69 Fleming, S. M., Ryu, J., Golfinos, J. G. & Blackmon, K. E. Domain-specific impairment in metacognitive accuracy following anterior prefrontal lesions. Brain: a journal of neurology 137, 2811–2822, doi:10.1093/brain/awu221 (2014).

70 Del Cul, A., Dehaene, S., Reyes, P., Bravo, E. & Slachevsky, A. Causal role of prefrontal cortex in the threshold for access to consciousness. Brain: a journal of neurology 132, 2531–2540, doi:10.1093/brain/awp111 (2009).

71 Maniscalco, B., Peters, M. A. K. & Lau, H. Heuristic use of perceptual evidence leads to dissociation between performance and metacognitive sensitivity. Atten Percept Psycho 78, 923–937, doi:10.3758/s13414-016-1059-x (2016).

72 McCurdy, L. Y. et al. Anatomical Coupling between Distinct Metacognitive Systems for Memory and Visual Perception. Journal of Neuroscience 33, 1897–1906, doi:10.1523/Jneurosci.1890-12.2013 (2013).

73 Griffin, D. & Tversky, A. The Weighing of Evidence and the Determinants of Confidence. Cognitive Psychol 24, 411–435, doi:DOI 10.1016/0010-0285(92)90013-R (1992).

74 Tversky, A. & Kahneman, D. Judgment under Uncertainty - Heuristics and Biases. Science 185, 1124–1131, doi:DOI 10.1126/science.185.4157.1124 (1974).

75 Zylberberg, A., Barttfeld, P. & Sigman, M. The construction of confidence in a perceptual decision. Front Integr Neurosci 6, 79, doi:10.3389/fnint.2012.00079 (2012).

76 Papadimitriou, C., White, R. L. & Snyder, L. H. Ghosts in the Machine II: Neural Correlates of Memory Interference from the Previous Trial. Cerebral cortex 27, 2513–2527, doi:10.1093/cercor/bhw106 (2017).

77 Hwang, E. J., Dahlen, J. E., Mukundan, M. & Komiyama, T. History-based action selection bias in posterior parietal cortex. Nature communications 8, doi:ARTN 1242 10.1038/s41467-017-01356-z (2017).

78 Akaishi, R., Umeda, K., Nagase, A. & Sakai, K. Autonomous Mechanism of Internal Choice Estimate Underlies Decision Inertia. Neuron 81, 195–206, doi:10.1016/j.neuron.2013.10.018 (2014).

79 Ratcliff, R. & McKoon, G. The diffusion decision model: Theory and data for two-choice decision tasks. Neural Comput 20, 873–922, doi:DOI 10.1162/neco.2008.12-06-420 (2008).

80 Fleming, S. M., Thomas, C. L. & Dolan, R. J. Overcoming status quo bias in the human brain. Proceedings of the National Academy of Sciences of the United States of America 107, 6005–6009, doi:10.1073/pnas.0910380107 (2010).

81 Bruno, M. A., Walker, E. A. & Abujudeh, H. H. Understanding and Confronting Our Mistakes: The Epidemiology of Error in Radiology and Strategies for Error Reduction. Radiographics 35, 1668–1676, doi:10.1148/rg.2015150023 (2015).

82 Prins, N. & Kingdom, F. A. A. Applying the Model-Comparison Approach to Test Specific Research Hypotheses in Psychophysical Research Using the Palamedes Toolbox. Frontiers in psychology 9, 1250, doi:10.3389/fpsyg.2018.01250 (2018).

83 Miller, G. A. Note on the bias of information estimates. Information Theory in Psychology: Problems and Methods, 95–100 (1955).

84 Lak, A. et al. Orbitofrontal cortex is required for optimal waiting based on decision confidence. Neuron 84, 190–201, doi:10.1016/j.neuron.2014.08.039 (2014).

85 Rouder, J. N., Speckman, P. L., Sun, D. C., Morey, R. D. & Iverson, G. Bayesian t tests for accepting and rejecting the null hypothesis. Psychon B Rev 16, 225–237, doi:10.3758/Pbr.16.2.225 (2009).

86 Wetzels, R. & Wagenmakers, E. J. A default Bayesian hypothesis test for correlations and partial correlations. Psychon B Rev 19, 1057–1064, doi:10.3758/s13423-012-0295-x (2012).

## Supplementary References

1 Urai, A. E., Braun, A. & Donner, T. H. Pupil-linked arousal is driven by decision uncertainty and alters serial choice bias. Nature communications 8, 14637, doi:10.1038/ncomms14637 (2017).

2 Braun, A., Urai, A. E. & Donner, T. H. Adaptive History Biases Result from Confidence-weighted Accumulation of Past Choices. The Journal of neuroscience: the official journal of the Society for Neuroscience, doi:10.1523/JNEUROSCI.2189-17.2017 (2018).

3 Suarez-Pinilla, M., Seth, A. K. & Roseboom, W. Serial dependence in the perception of visual variance. J Vision 18, doi:ARTN 4 10.1167/18.7.4 (2018).

4 Samaha, J., Switzky, M. & Postle, B. R. Confidence boosts serial dependence in orientation estimation. J Vision 19, doi:ARTN 25 10.1167/19.4.25 (2019).

5 Desender, K., Boldt, A. & Yeung, N. Subjective Confidence Predicts Information Seeking in Decision Making. Psychological science 29, 761–778, doi:10.1177/0956797617744771 (2018).

6 Gherman, S. & Philiastides, M. G. Neural representations of confidence emerge from the process of decision formation during perceptual choices. NeuroImage 106, 134–143, doi:10.1016/j.neuroimage.2014.11.036 (2015).

7 Murphy, P. R., Robertson, I. H., Harty, S. & O’Connell, R. G. Neural evidence accumulation persists after choice to inform metacognitive judgments. eLife 4, doi:ARTN e11946 10.7554/eLife.11946 (2015).

